# Mapping embryonic mouse lung development using enhanced spatial transcriptomics

**DOI:** 10.1101/2025.11.22.688293

**Authors:** Pengfei Zhang, Benjamin K. Law, Katharine Goodwin, Michelle M. Chan, Celeste M. Nelson

## Abstract

The mature mammalian lung contains ∼50 different cell types that are located in precise positions along and around the airway epithelial tree. Here, we investigated how these spatial patterns are progressively established during development. Specifically, we optimized a microfluidics-enabled, deterministic barcoding-based sequencing (DBiT-seq) workflow to achieve highly sensitive spatial mapping of tissue sections from early embryonic mouse lungs. Spatial mapping of epithelial populations revealed that *Sox9*^+^ epithelial progenitors are located throughout the epithelial tree at early stages before becoming confined to the distal ends. Spatial mapping of mesenchymal populations revealed a previously unrecognized mesenchymal cluster that expresses high levels of extracellular matrix proteins and appears to promote embryonic lung innervation. These findings provide new insights into the spatial organization and cellular dynamics underlying early lung development, and demonstrate the power of spatial transcriptomics to uncover hidden patterns and populations of cells.

## Introduction

The spatial architecture of the developing lung plays a key role in its morphogenesis. Initiated at embryonic day (*E*) 9.5, the mouse lung buds from the ventral foregut endoderm and then recursively branches to form a tree-shaped structure^1,2^. As the epithelium branches, the surrounding mesenchyme differentiates into diverse lineages that are localized to distinct regions throughout the lung. These include mesothelium enveloping the outer surface of the lung, cartilage strengthening the trachea, smooth muscle wrapping around conducting airways, and pericytes supporting capillary endothelium. Collectively, the mesenchymal populations provide inductive signals that help specify proximodistal epithelial patterning, establishing *Sox2*^+^ epithelium in proximal airways and *Sox9*^+^ epithelial progenitors at the distal tips^1–3^.

An outstanding question in the field is the precise location of multipotent epithelial progenitors and how their location shifts during the establishment of the proximodistal axis in the developing lung epithelium^4^. Lineage-tracing studies suggest that multipotent *Id2*^+^ epithelial cells, located at the distal tips of the tree, contribute to both the proximal conducting airways and distal alveoli depending on the developmental stage^5^. Subsequent work showed that two nested waves – marked by the expression of SOX2 and SOX9 – demarcate a compartment boundary between proximal conducting airways and distal gas-exchange epithelium^4^. However, the precise spatiotemporal distribution of multipotent progenitors, especially at early embryonic stages, remains largely undefined beyond insights from immunostaining or in situ hybridization of a limited set of targets^4–7^.

In parallel, a major challenge in studying lung mesenchyme has been to classify this highly heterogeneous tissue into different subtypes, particularly the mesenchymal progenitors that lack unique gene markers. Early studies showed that *Wnt2*^+^ mesenchymal progenitor cells give rise to multiple mesenchymal lineages during embryonic development, including airway smooth muscle^8^. Yet, subsequent efforts to define smooth-muscle progenitors revealed no unique molecular markers^9^. Clonal labeling experiments found that smooth-muscle progenitors are spatially restricted to the mesenchyme positioned ahead of the branching epithelium, underscoring the importance of anatomical location in subtype specification^10^. Consistently, recent studies identified anatomically distinct mesenchymal subtypes that serve different roles in response to tissue injury and alveolar epithelial cell proliferation^11,12^. Collectively, these findings highlight that spatial context is a key determinant of mesenchymal identity and function.

Incorporating anatomical location into classification frameworks may therefore help define mesenchymal subtypes and elucidate their roles in lung development.

Spatial transcriptomics offers the opportunity to complement single-cell analysis and provide spatial context to the different cell types within a developing tissue, even those that are difficult to isolate into single cells for single-cell RNA sequencing (scRNA-seq), such as the epithelium^13,14^. Consistently, efforts have recently been made to obtain comprehensive spatiotemporal maps of the embryonic lung^15–18^ . Unfortunately, current whole-transcriptomic spatial-mapping approaches suffer from limited resolution and/or sensitivity, hindering spatial mapping at the single-cell level with high transcript detection sensitivity. Emerging technologies, such as Visium HD^19^, Slide-seqV2^20^, Seq-Scope^21^, Stereo-seq^22^, and OpenST^23^, achieve single-cell resolution but the detection sensitivity remains low due to inefficient mRNA diffusion and capture^23,24^. In contrast, microfluidics-based approaches, such as deterministic barcoding-based sequencing (DBiT-seq), enhance detection sensitivity by delivering spatial DNA barcodes directly into the tissue, bypassing barriers to mRNA diffusion^25^ while achieving close-to-single-cell resolution. Nevertheless, the transcript recovery for microfluidics-based approaches remains low compared to scRNA-seq^23,24^.

Here, we improved the detection sensitivity of single-cell spatial transcriptomics by optimizing the DBiT-seq workflow^25^. To highlight the benefits of this optimized workflow, we generated spatial datasets for tissue sections of the embryonic mouse lung at close-to-single-cell resolution. Combining our spatial transcriptomic data with reference scRNA-seq allowed us to spatially map known cell types and uncover previously undescribed cell populations and tissue-tissue interactions. We confirmed that proximal and distal epithelial cells exhibit distinct patterns of gene expression, but our spatial mapping unexpectedly revealed that ‘distal’ epithelial progenitor cells are located in proximal regions of the lung at early stages of development. Moreover, we identified the spatial locations of three subclusters of *Wnt2*^+^ mesenchymal progenitors that show distinct spatiotemporal patterns over time. Finally, we discovered a novel mesenchymal cell type that expresses an abundance of extracellular matrix (ECM) markers and related genes, which we refer to as “Matrix Mesenchyme”. Our spatial transcriptomic mapping and cell-cell interaction analyses reveal that Matrix Mesenchyme localizes near the trachea and medial boundaries of the lung lobes, where innervation of the embryonic lung typically starts^26,27^, and interacts with neurons to promote innervation of the developing organ. Together, these findings establish our optimized workflow and datasets as valuable resources for the community, offering new insights into the spatial organization and cellular dynamics of early lung development.

## Results

### Early embryonic lung development profiled using an optimized spatial transcriptomics workflow

To investigate the spatial dynamics of the embryonic mouse lung, we optimized a microfluidics-based DBiT-seq workflow to enable unbiased, high-resolution spatial mapping of the transcriptome (**Fig. 1a, Fig. S1-3**). DBiT-seq uses microfluidic chips to deliver oligonucleotides into tissue sections for spatial barcoding at a resolution that corresponds to the width of the channels on the chip (10-50 μm). Building upon a published DBiT-seq workflow^16^, we increased the capture efficiency by optimizing tissue fixation^28^, adding a randomly primed second-strand synthesis step^20,29^, and employing a template-switch step in the *in situ* reverse-transcription phase (**Fig. S1, Fig. S2a-g**). This optimized DBiT-seq protocol yielded a 55-150% increase in UMI detection per pixel when tested on sections of whole embryos using 25-μm-resolution chips, after standardizing for pixel size and sequencing depth (**Fig. S2h**). Our optimized workflow also recovers more UMIs (∼1,000-1,250 UMI/100 μm^2^) than the original DBiT-seq protocol (∼600-1,200 UMI/100 μm^2^) and the recently established Open-ST (800 UMI/100 μm^2^) when tested on sections of whole embryos using 10-μm-resolution chips (**Fig. 1b**)^23^. We therefore used this approach to spatially map the developing mouse lung.

**Fig. 1.**
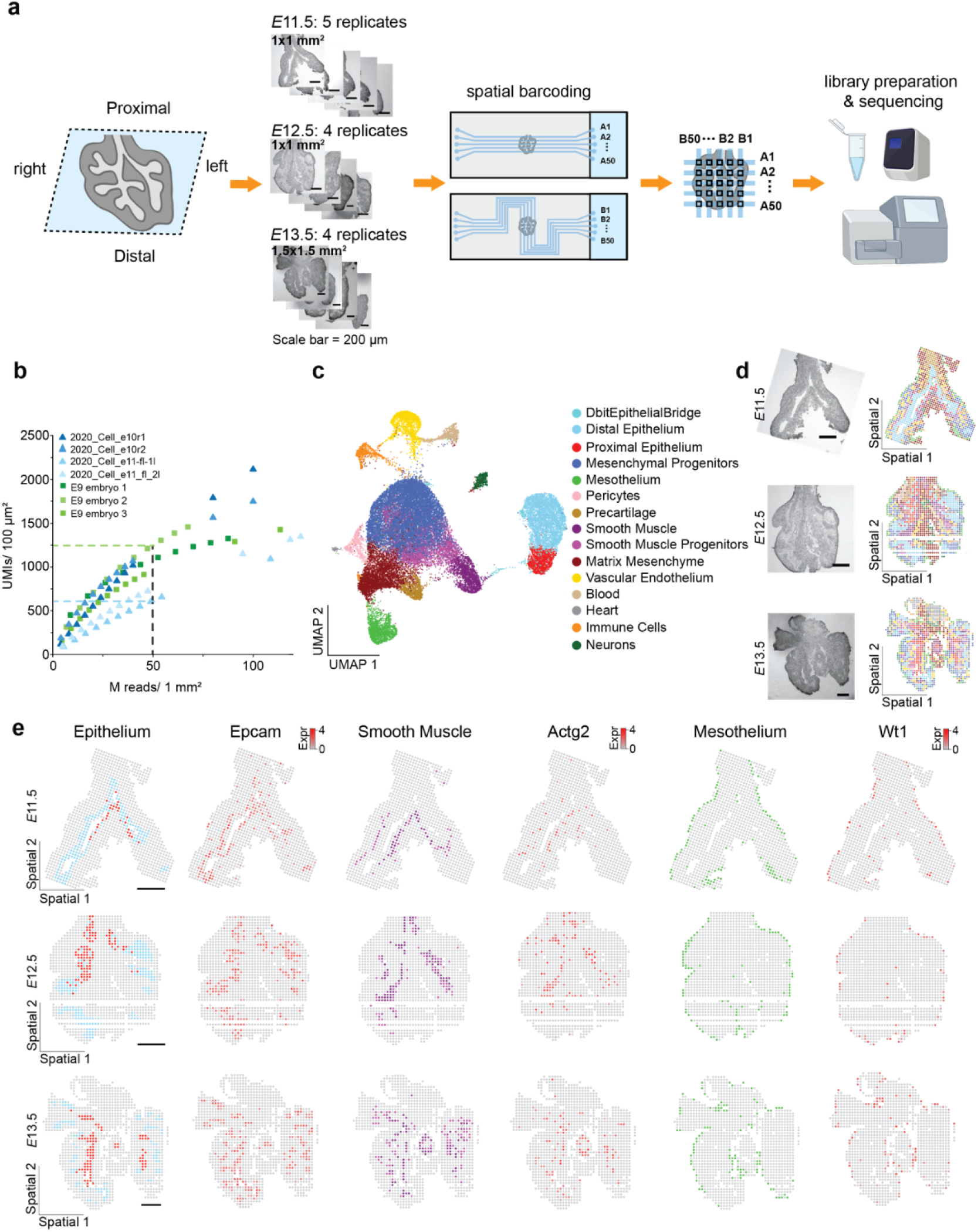
Spatial mapping of cell types in the embryonic mouse lung. (**a**) Schematic overview of workflow for spatial transcriptomic mapping of the embryonic mouse lung. (**b**) Saturation curves of the optimized DBiT-seq protocol (green) as compared to the original, published DBiT-seq protocol (blue) for sections of whole embryos analyzed using 10-μm-resolution chips. (**c**) UMAP showing cell-type clustering for the embryonic mouse lung. (**d**) Brightfield images of representative sections and color-coded maps of all cell types at each stage of development. Scale bars, 200 µm. (**e**) Color-coded maps of epithelium, smooth muscle, and mesothelium, as well as their top gene markers at each stage. Scale bars, 200 µm.

We focused on coronal sections of embryonic lungs acquired from *E*11.5-*E*13.5, when the epithelial tree is established (**Fig. 1a, Fig. S3a**). Specifically, we obtained 13 total replicates: 5 at *E*11.5 and 4 each at *E*12.5 and *E*13.5. The *E*11.5 and *E*12.5 replicates were analyzed at 10-μm resolution, while the *E*13.5 replicates were analyzed at 15-μm resolution to ensure we covered the entire tissue section. In total, we recovered 15,037 high-quality pixels with a median value of 1,943 UMIs and 1,251 genes per replicate, consistent with tests using whole embryos (**Fig. 1b, Fig. S2i, Fig. S4a-b**). We observed a slightly higher recovery (2,221 UMIs and 1,480 genes per pixel, respectively) at *E*13.5, likely due to the larger pixel size.

To annotate cell identities, we first integrated our spatial transcriptomics data with published scRNA-seq datasets^30–33^ and clustered the integrated data using the Louvain algorithm (see **Methods**)^34^. Clusters, which contain both single cells and spatial pixels, were then annotated according to canonical marker genes, identifying major cell types such as Proximal and Distal Epithelium, Smooth Muscle, Mesothelium, and Mesenchymal Progenitors (**Fig. 1c, Fig. S4c**). The proportions of annotated cell types are consistent within replicates of the scRNA-seq and spatial transcriptomic data, respectively. However, we noticed that our spatial transcriptomics dataset captured a higher proportion of epithelial cells than the scRNA-seq dataset (**Fig. S3c**). This difference might be attributed to the fact that the latter approach requires dissociation of the organ into single cells, which has been shown to be challenging for epithelial tissues, including that of the embryonic lung^13,14^.

Mapping of the annotated cell types onto the tissue sections showed the expected locations, with an *Epcam*^+^ epithelial tube surrounded by *Actg2*^+^ smooth muscle cells and a peripheral layer of *Wt1*^+^ mesothelium (**Fig. 1d-e**), consistent across replicates and timepoints (**Fig. S5a-c**). To determine whether the optimized protocol is close to resolving single cells per pixel, we carried out conditional autoregressive-based deconvolution (CARD)^35^. This analysis generated similar spatial maps of the major cell types of the embryonic mouse lung (**Fig. S5d-e**). However, we also found a portion of pixels that presented as a mixture of different cell types. For example, 21.5% of pixels annotated as epithelium were predicted to contain <50% epithelial cells. The predicted pixel proportion containing mixed cell types is consistent with the challenge of guaranteeing single-cell capture for spatial transcriptomics mapping^36^. Nevertheless, our optimized DBiT-seq workflow enables high-resolution (10-µm) mapping of the major cell types of the early embryonic mouse lung with high sensitivity. To avoid any confounding issues from the possibility of cellular mixtures in spatial data, all differential gene-expression analyses in the following sections were conducted using scRNA-seq data.

### Spatiotemporal mapping of the epithelium reveals unexpected pattern of *Sox9*^+^ progenitors

We took advantage of this dataset to investigate the differentiation trajectories and spatiotemporal distributions of the major tissues in the embryonic mouse lung. We first focused on the epithelium. According to previous lineage-tracing studies, the lung epithelium is initially patterned along its proximodistal axis, with the distal *Sox9*^+^*Id2*^+^ epithelial cells serving as the progenitors for different mature epithelial cell types (**Fig. 2a**)^1,2,37^. While these *Sox9^+^Id2*^+^ epithelial progenitors have been shown to localize to the distal regions of the epithelial tree from *E*11.5, it is not clear if they are spatially restricted or when that restriction may occur^5^. As a result, the precise spatiotemporal distribution of these progenitors, especially at early embryonic stages, remains largely undefined.^4–7^

**Fig. 2.**
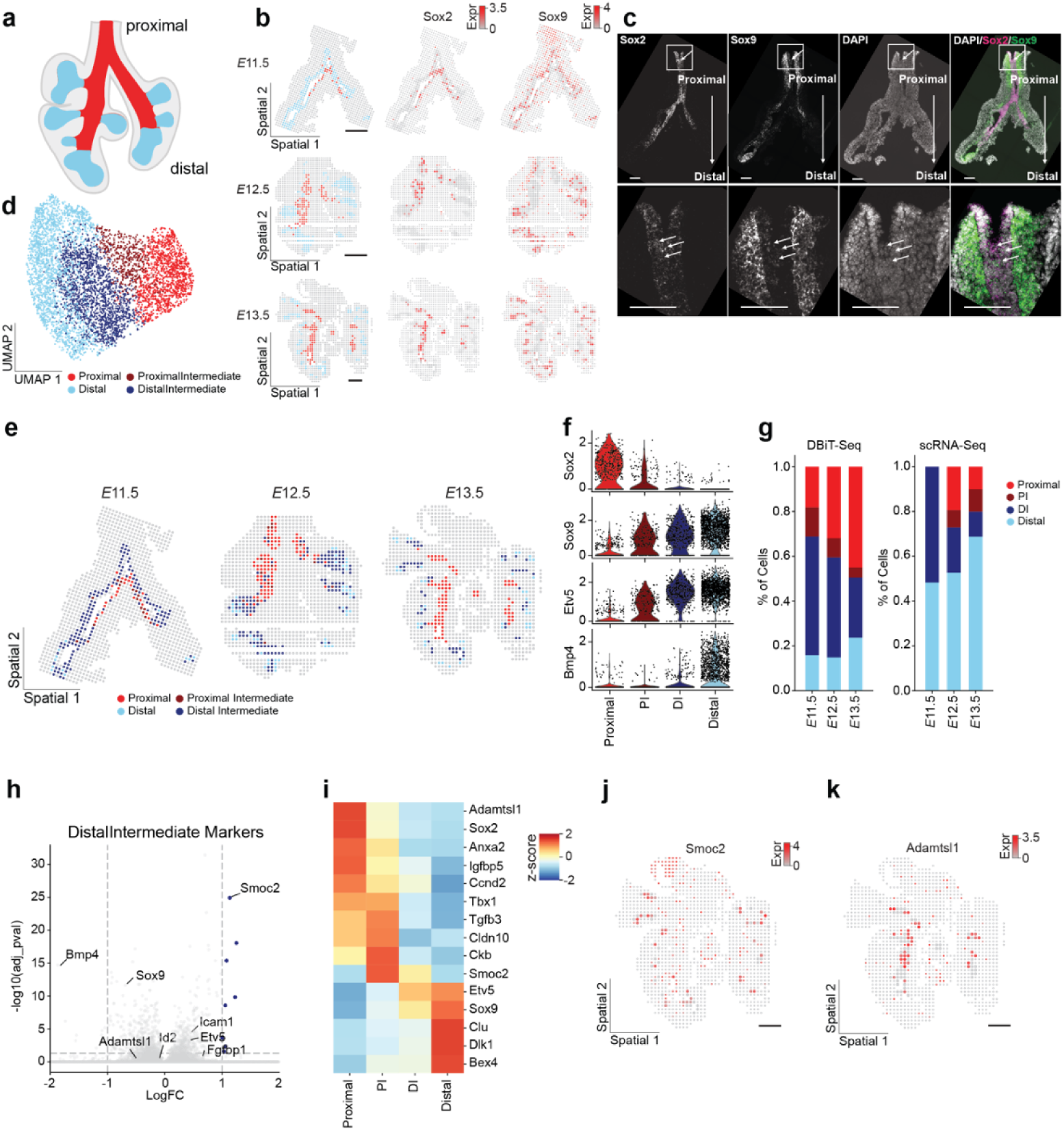
Spatiotemporal mapping of the epithelium reveals unexpected pattern of multipotent progenitors. (**a**) Schematic of current conceptual model of lung epithelial development. (**b**) Color-coded maps of proximal and distal epithelium and their top gene markers at each stage, including *Sox2* and *Sox9*. Scale bars, 200 µm. For ease of visualization, larger and darker pixels are used to highlight epithelial cells. (**c**) Fluorescence images of hybridization analysis for DAPI, *Sox2*, and *Sox9*. Scale bars, 100 µm. The raw images were rotated and cropped to orient the trachea upward, and the empty space was filled with a rectangular black box. (**d**) UMAP of epithelial subclustering reveals 4 different subtypes. (**e**) Color-coded maps of epithelial subclusters from (d). (**f**) Violin plots of *Sox2*, *Sox9*, *Etv5*, and *Bmp4*. (**g**) Bar plots of epithelial subcluster compositions for DBiT-seq and scRNA-seq. (**h**) Volcano plot of differential gene-expression analysis between Distal Intermediate (DI) and the other epithelial cell subtypes. (**i**) Heatmap of the differentially expressed genes. (**j-k**) Color-coded maps of *Smoc2* and *Adamtsl1* at *E*13.5. For ease of visualization, larger and darker pixels are used to highlight PI/DI and Proximal epithelial cells. Scale bars, 200 µm.

Our spatial mapping provides a comprehensive delineation of the spatiotemporal pattern within the epithelium. Specifically, we identified two distinct epithelial clusters that express high levels of the proximal and distal epithelial markers, *Sox2* and *Sox9*, respectively (**Fig. 2b**). However, the spatial mapping of these clusters reveals that the *Sox9*^+^ ‘distal’ epithelial cells, annotated using additional markers including *Etv5* and *Id2*, are located in both the proximal and distal regions of the epithelium at *E*11.5 (**Fig. 2b, Fig. S6-8**). Nonetheless, we observe the expected distal localization of the *Sox9*^+^ cluster at *E*12.5 and *E*13.5 (**Fig. 2b, Fig. S6-8**). To validate the spatial position of the proximal and distal epithelial clusters, we profiled the expression of *Sox2*, *Sox9*, and *Shh* (a marker of the entire epithelium) in lung sections using fluorescence in situ hybridization. Our results confirmed that the proximal epithelium contains *Sox9^+^*epithelial cells at *E*11.5, which gradually diminish from this region at later time points (**Fig. 2c, Fig. S9**).

Quantifying the levels of *Sox2* and *Sox9* within *Shh*^+^ epithelial pixels revealed that *Sox9* is expressed in a subset of *Sox2^+^* pixels, the fraction of which decreases from *E*11.5 to *E*13.5 (**Fig. S9, Fig. S10a**). Analysis of scRNA-seq reference data also uncovered a small population (6-10%) of *Sox2*^+^*Sox9*^+^ cells from *E*11.5 to *E*13.5 (**Fig. S10b**). *Sox2*^+^*Sox9*^+^ epithelial cells have been previously reported in the embryonic human lung but not in mouse, to our knowledge^38^.

This discrepancy may reflect differences in sampling strategies, as prior studies largely concentrated on the distal epithelium and later developmental stages^6,7,39^.

We next subclustered the epithelial cell populations to identify possible transition states between the *Sox2*^+^ and *Sox9*^+^ cells (**Fig. 2d-e, Fig. S11a-c**). This analysis revealed spatially distinct epithelial subclusters, including Proximal, Proximal Intermediate (PI), Distal Intermediate (DI), and Distal, located along the proximodistal axis of the epithelial tube (**Fig. 2e, Fig. S11a-c**).

These subclusters show gradients of *Sox2* and *Sox9* markers as well as other markers including *Igfbp5*, *Ccnd2*, *Cdnk1c*, *Etv5*, *Id2*, *Bmp4* and *Sftpc* (**Fig. 2f, Fig. S11d**). There is a shift in the composition of these subclusters over time: PI and DI states are observed primarily at *E*11.5 and transition to proximal and distal epithelial cells at *E*12.5 and *E*13.5 (**Fig. 2g**).

To identify possible markers involved in the initial commitment towards the proximal or distal fates, we analyzed the genes that are highly expressed in both PI and DI (**Fig. 2h-i, Fig. S12a-c**). We found that *Smoc2* is upregulated in both PI and DI compared to the other subclusters (**Fig. 2h-i, Fig. S12a-c**) and spatial maps showed that *Smoc2* is specifically expressed at high levels in the epithelium immediately adjacent to the proximal- and distal-most ends of the epithelial tubes (**Fig. 2j**). *Adamtsl1*, upregulated in Proximal Epithelium (**Fig. 2i**), is localized to the proximal side (**Fig. 2k, Fig. S12c**), while *Bmp4*, *Adamts18*, and *Thbd*, which are highly expressed in Distal Epithelium (**Fig. 2f, Fig. S12c**), are at the distal tips (**Fig. S13**). *Bmp4*, *Adamts18*, and *Etv4* have been previously shown to be critical for distal alveolar differentiation^40–42^, but the exact function of *Smoc2* and *Adamtsl1* in epithelial differentiation requires further investigation. Notably, *Icam1* and *Fgfbp1*, which have been used as markers for intermediate epithelium, were either not identified as a specific marker (*Icam1*) or barely enriched in our analysis (*Fgfbp1*) (**Fig. 2h, Fig. S12a-b, and Fig. S12d**). We also built a transcriptional pseudotime axis that showed the progression between Proximal and Distal epithelial subclusters (**Fig. S12e-g**). The regression analysis and module-enrichment analysis along the pseudotime axis revealed similar gene-expression enrichment in each state (**Fig. S12h**). Altogether, our spatial mapping highlights spatiotemporal dynamics of epithelial transitional states and suggests possible markers of the intermediate states between proximal and distal epithelium.

### Spatial mapping reveals distinct locations of mesenchymal progenitors and provides spatial context for smooth-muscle differentiation

To study the highly heterogenous embryonic lung mesenchyme, we isolated the mesenchymal cell types and subclustered the dataset. This analysis revealed eight subclusters, including both mesenchymal progenitors and mature mesenchymal states (**Fig. 3a-c, Fig. S14a**). We first focused on the three progenitor states that express high levels of *Wnt2*: Mesenchymal Progenitor 1 (MP1) is in the periphery of the lung lobes and observed at all time points; Mesenchymal Progenitor 2 (MP2) is in the periphery of the lobes as well but first appears at *E*12.5; and Mesenchymal Progenitor 3 (MP3) is close to the vascular endothelium and appears at *E*12.5 and *E*13.5 (**Fig. 3d, Fig. S15a-c, and Fig. S15g-i**). The spatial maps of these Mesenchymal Progenitor populations match the spatial patterns of markers for mesenchymal progenitors, including *Wnt2*, *Hoxa5*, and *Hoxb5* (**Fig. 3d, Fig. S16a-c**), and are consistent with recent maps of mesenchymal progenitors in the embryonic human lung ^18,43,44^ . We performed differentially expressed gene (DEG) analysis to identify markers for each subcluster, which showed that MP1 is enriched in genes associated with cell-cycle regulation and cell division (**Fig. S14a-b**). In contrast, MP2 and MP3 express lower levels of proliferation markers and high levels of *Rspo2/Fgf10* and *Tagln2/Angpt1*, respectively (**Fig. S14c-d**). We also observed that MP2 expresses high levels of *Hoxb8*, *Calb2*, and *Asz1*, which exhibit a unique spatiotemporal pattern at the periphery of the lung (**Fig. S14c, Fig. S16a-c**). The specific functions of these genes in lung development remain to be further investigated.

**Fig. 3.**
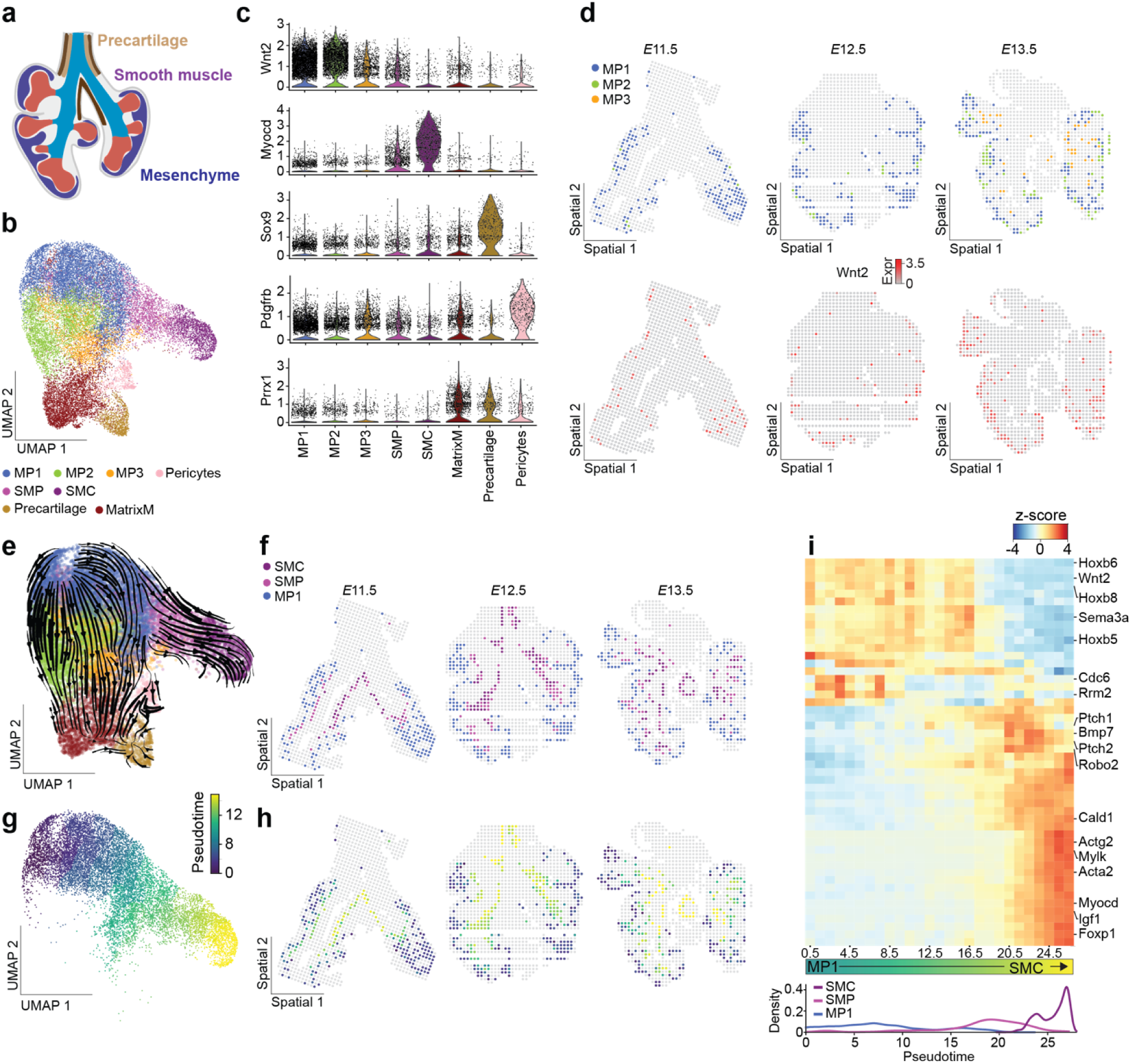
Distinct localization of mesenchymal progenitors and spatial context for smooth muscle differentiation in embryonic lung. (**a**) Schematic of mesenchyme within embryonic mouse lung. (**b**) UMAP of mesenchymal subclusters. (**c**) Violin plots of top gene markers for each mesenchymal subcluster. (**d**) Color-coded maps of mesenchymal progenitors at each stage of lung development, together with spatial maps of the marker *Wnt2*. (**e**) UMAP of RNA velocity for mesenchymal populations. (**f**) Color-coded maps of Mesenchymal Progenitor 1 (MP1), Smooth Muscle Progenitors (SMP), and Smooth Muscle Cells (SMC). (**g**) UMAP of pseudotime inference for the smooth-muscle trajectory. (**h**) Spatial pseudotime mapping of smooth-muscle trajectory at each stage of lung development. (**i**) Heatmap of gene markers along the pseudotime trajectory and graph of density of cell states along the pseudotime from (g).

The intermediate and mature cell states in our dataset include Smooth Muscle Progenitors (SMP; *Wnt2*^low^*Myocd*^low^*Ptch1*^high^) and Smooth Muscle (SMC; *Myocd*^high^), both surrounding the epithelium, Precartilage (*Sox9*^high^) surrounding the trachea and upper airways, and Pericytes (*Pdgfrb*^high^) adjacent to vascular endothelial cells (**Fig. 3c**, **Fig. S14a, and Fig. S15d-f**).

Additionally, we identified a population that we label as Matrix Mesenchyme (MatrixM; *Prrx1*^high^*Ptn*^high^*Col1a1*^high^) due to its high expression of ECM proteins and related factors (**Fig. 3b-c, Fig. S14a**).

Given the presence of both progenitor and mature smooth-muscle-cell states in our atlas, we proceeded to examine the spatial dynamics of mesenchymal progenitors as they differentiate into smooth muscle cells, a process that has been well described using clonal lineage tracing and scRNA-seq^9,10,18,30,31,45,46^. We reasoned that MP1 is the progenitor source for all mesenchymal cells due to its prevalence at *E*11.5 compared to MP2 and MP3. RNA velocity, which uses the ratios of unspliced/spliced transcripts in RNA-seq data to inform trajectory direction, also infers that MP1 is the source for mesenchymal trajectories (**Fig. 3e**). We then subset the data to include the subclusters of interest and constructed a pseudotime trajectory starting from MP1, through Smooth Muscle Progenitors, and into Smooth Muscle (**Fig. 3f-h**). We performed regression and module analysis to identify transcriptional modules that vary along the pseudotime trajectory (**Fig. 3i**). This analysis identified distinct modules that contain genes specifically expressed in MP1, Smooth Muscle Progenitor, and Smooth Muscle (**Fig. 3i**). As expected, we observed high expression of markers of mesenchymal progenitors (*Wnt2* and *Hoxb5*) and mature smooth muscle (*Actg2*, *Mylk*, *Myl9* and *Mycod*) at the beginning and end of the trajectory, respectively. Consistently, as compared to MP1, we found that Smooth Muscle cells downregulate expression of the mesenchymal progenitor marker *Wnt2*, and upregulate expression of smooth-muscle markers, as expected (**Fig. S16d**). Notably, our analysis also revealed that Smooth Muscle Progenitors upregulate *Ptch1* and *Ptch2*, receptors for Shh, and that the expression of *Ptch1* drops in Smooth Muscle (**Fig. 3i, Fig. S16e**). These observations are consistent with previous reports that Shh expressed by the airway epithelium induces the surrounding mesenchyme to differentiate into airway smooth muscle^47^.

To test whether our dataset could provide spatial context for the differentiation trajectories of the mesenchymal subclusters, we mapped the pseudotime analysis spatially. We observed a spatial association between smooth muscle maturation and lung topology: the most immature MP1 localizes to the periphery of the lung, while Smooth Muscle Progenitors and Smooth Muscle cells progressively align closer to the bronchi along the distal-to-proximal axis (**Fig. 3f-h**). This spatial pattern aligns with previously published observations that mesenchymal progenitors adjacent to epithelial tips are recruited to the sites of branching to differentiate into smooth muscle^10,48^. This spatial pattern is also consistent with that reported for the embryonic human lung^18^, but with higher resolution. Collectively, integrating spatial location allowed us to map distinct mesenchymal subtypes and provides spatial context for the smooth-muscle differentiation trajectory.

### Matrix Mesenchyme localizes near Neurons and enriches genes driving innervation

We next focused on the Matrix Mesenchyme population, which expresses high levels of several ECM-related genes, including *Col1a1*, *Tmtc2*, *Col1a2*, and *Col3a1*, as well as markers *Ptn*, *Igfbp5*, *Rspo2*, *Prrx1*, and *Osr1* (**Fig. 4a**, **Fig. 4c, and Fig. S17a-c**). Notably, previous studies identified this population of cells but often lumped it together with mesenchymal progenitors (**Fig. S3c**)^30,32^. While it shares some genes with mesenchymal progenitors, such as those related to collagen, Matrix Mesenchyme expresses minimal levels of the mesenchymal progenitor marker, *Wnt2* (**Fig. 3c, Fig. S14a**)^8,32,43^. Our spatial dataset revealed that Matrix Mesenchyme is consistently located around the trachea and along the medial boundaries of the lobes, distinct from *Wnt2*^high^ Mesenchymal Progenitors that localize to the lung periphery (**Fig. S17a-c, Fig. S18a-c**). To verify the spatial location of these cells, we conducted fluorescence in situ hybridization analysis for *Prrx1*, *Osr1*, and *Sox9*. We used the latter transcript to distinguish between Precartilage (*Sox9*^high^) and Matrix Mesenchyme (*Sox9*^low^), as Precartilage also expresses both *Prrx1* and a low level of *Osr1*, and localizes to the trachea (**Fig. 3c, Fig. S18d**). We observed a spatial pattern of *Prrx1*^high^*Osr1*^high^*Sox9*^low^ cells, consistent with our spatial dataset (**Fig. 4b**). We therefore conclude that Matrix Mesenchyme is a distinct population of cells in the embryonic mouse lung and likely does not function as a mesenchymal progenitor.

**Fig. 4.**
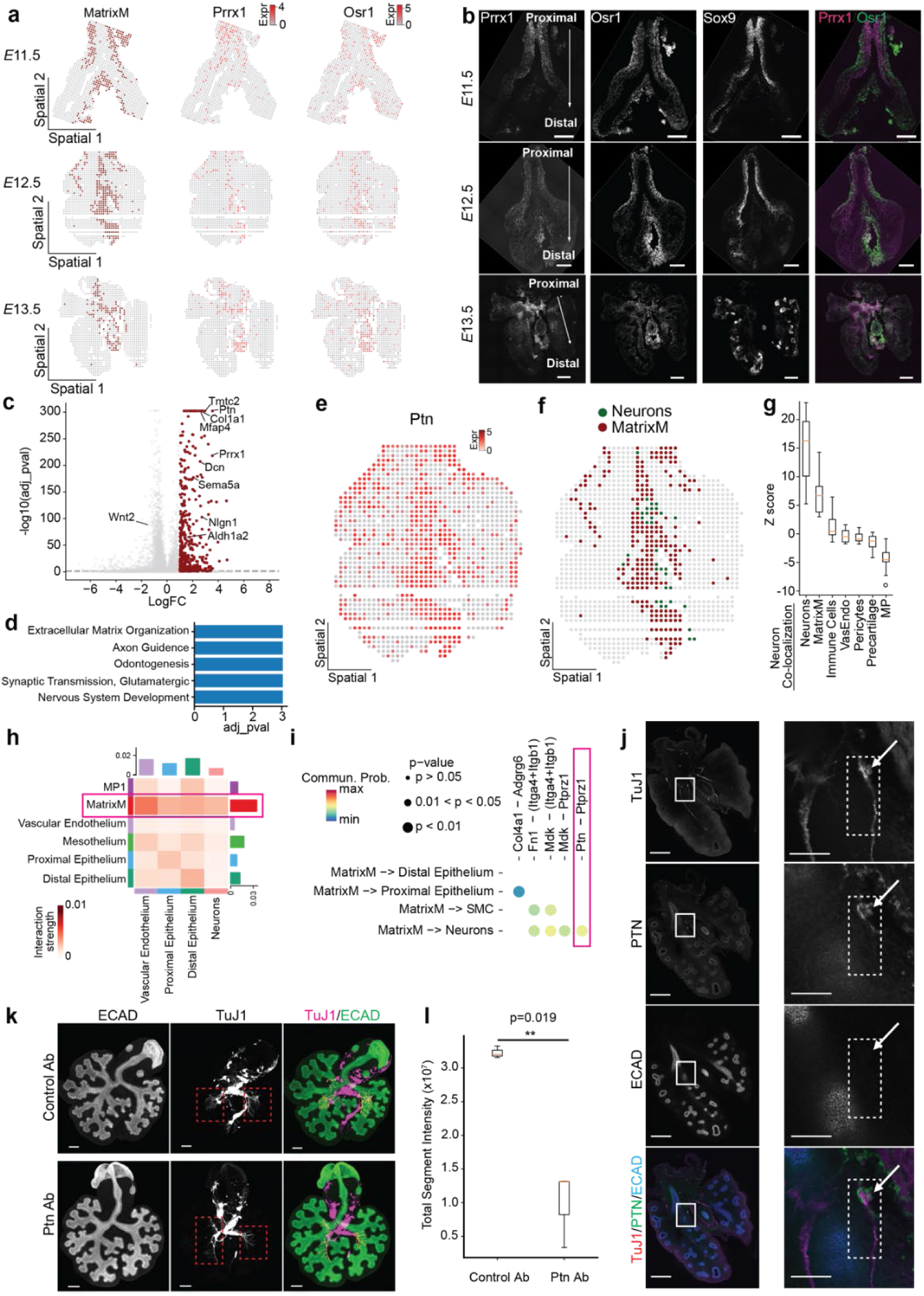
Spatial profiling of Matrix Mesenchyme reveals ECM enrichement and roles in lung innervation. (**a**) Color-coded maps of Matrix Mesenchyme (MatrixM), together with spatial maps of the markers *Prrx1* and *Osr1* at each stage. (**b**) Fluorescence in situ hybridization analysis of *Prrx1* (green), *Osr1* (magenta), and *Sox9*. Scale bars, 200 µm. The raw images were rotated and cropped to orient the trachea upward, and the empty space was filled with a rectangular black box. (**c**) Volcano plot of differential gene-expression analysis between Matrix Mesenchyme and the other cell types. (**d**) Gene Ontology analysis for the top-enriched genes in Matrix Mesenchyme. (**e**) Color-coded spatial map of *Ptn* expression at *E*12.5. (**f**) Color-coded map of Neurons and Matrix Mesenchyme. (**g**) Z-score plot for neighborhood-enrichment analysis of Neurons. MatrixM: Matrix Mesenchyme; VasEndo: Vascular Endothelium; MP: Mesenchymal Progenitors. (**h**) Heatmap of communication probability between different cell types. (**i**) Top ligand-receptor interactions between Matrix Mesenchyme and other cell types. SMC: Smooth Muscle Cells. (**j**) Immunofluorescence analysis for TuJ1, PTN, and E-cadherin for a whole lung section (left) and zoom-in view (right). Scale bars, 200 µm (left), 50 µm (right). (**k**) Immunofluorescence analysis for E-cadherin and TuJ1 in lung explants cultured in the presence of anti-PTN or control antibodies. Scale bars, 100 µm. (**l**) Graph showing quantification of innervation in lung explants. N = 3.

To investigate the potential function of Matrix Mesenchyme, we performed Gene Ontology (GO) analysis on the most upregulated genes in this population compared to all other mesenchymal cells (**Fig. 4c-d**). As expected from its high expression of ECM-related genes, Matrix Mesenchyme shows strong enrichment in gene sets associated with Extracellular Matrix Organization (**Fig. 4d**, p =9.3×10^-4^). Matrix Mesenchyme is also enriched in innervation-related terms, including Axon Guidance (p = 9.3×10^-4^), Synaptic Transmission, Glutamatergic (p = 4.7×10^-6^), Nervous System Development (p = 5.1×10^-6^), and Neuron Projection Guidance (p = 6.0×10^-6^), suggesting a potential function in development of the nervous system (**Fig. 4d**). This enrichment is driven in part by high expression of genes including *Ptn* among others (**Fig. 4e**). *Ptn* encodes for pleiotrophin (PTN), a multifunctional growth factor that promotes the migration and neurite outgrowth of embryonic neurons^49,50^. Consistently, two PTN receptors, *Sdc3* and *Ptprz1*, are both specifically expressed in Neurons at these stages of lung development (**Fig. S18e**). To further investigate the relationship between Matrix Mesenchyme and Neurons, we performed neighborhood-enrichment analysis and found that Matrix Mesenchyme and Neurons are located in close proximity to each other (**Fig. 4f-g, Fig. S18a-c**).

Consistent with their spatial patterns, cellchatV2^51,52^ revealed significant ligand-receptor interactions between Matrix Mesenchyme and Neurons (**Fig. 4h-i**), with one notable example being the PTN-PTPRZ1 ligand-receptor axis (**Fig. 4i**). To map PTN signaling and neurons, we conducted immunofluorescence analysis for PTN together with neuronal (TuJ1) and epithelial (E-cadherin) markers. This analysis revealed that PTN-expressing cells are located adjacent to neurons but not epithelial cells (**Fig. 4j, Fig. S19**). To determine the role of PTN in neuronal patterning, we cultured embryonic lung explants in the presence of a PTN function-blocking monoclonal antibody, which led to a significant reduction in the extent of innervation in the lung (**Fig. 4k-l, Fig. S20**, p = 0.019). These data suggest that PTN serves as a signal to promote innervation of the early embryonic mouse lung. Our spatial mapping thus identifies Matrix Mesenchyme as a novel mesenchymal subcluster with distinct gene expression and spatial locations, which may promote lung innervation during early stages of lung development.

## Discussion

Here, we optimized DBiT-seq and demonstrated improved transcript recovery compared to existing whole-transcriptomic spatial transcriptomics methods that rely on mRNA diffusion. Microfluidics-based approaches such as DBiT-seq enhance detection sensitivity by delivering spatial DNA barcodes directly into the tissue, bypassing the need for mRNA to diffuse to capture probes^25^. Our refinement of tissue-section fixation and addition of random-priming-assisted second-strand synthesis further improved detection sensitivity, thus enabling capture of more UMIs and higher gene counts.

We tested whether our optimized spatial transcriptomics protocol could uncover the spatial dynamics of key cell states by using the embryonic mouse lung as a model. Our spatial analysis confirms epithelial differentiation along the proximodistal axis of the organ but surprisingly reveals that *Sox9*^+^*Etv5*^+^*Id2*^+^ epithelial progenitors are localized in proximal regions of the epithelial tree at *E*11.5. This unexpected spatial pattern contrasts with the prevailing conceptual model of lung development, which assumes an initial proximodistal patterning of the epithelial tree, with distal epithelial cells serving as the sole progenitors for different mature epithelial cell types^1,2,5^. Nevertheless, our findings reconcile previous lineage-tracing studies showing that *Id2*^+^ progenitors, located proximally at *E*12.5 in our spatial data (**Fig. S7d-f**), give rise to both bronchiolar and alveolar cell types when labeled before *E*12.5 but only alveolar cell types when labeled at *E*16.5^5^. Our findings suggest that additional studies using precisely time-controlled lineage tracing are needed to validate the differentiation trajectories of the epithelium at different time points.

Our optimized pipeline was especially useful for disentangling the heterogeneous mesenchymal compartment. We uncovered a previously undescribed mesenchymal cluster, Matrix Mesenchyme, which localizes near the trachea and medial boundaries of the lobes and is characterized by high expression of ECM-related proteins. While this cluster is present in scRNA-seq reference data, it was previously classified as part of the mesenchymal progenitor population^30,32^. Two lines of evidence support our conclusion that Matrix Mesenchyme represents a distinct cell type. First, Matrix Mesenchyme expresses low levels of *Wnt2* and markers of proliferation, which are expressed at high levels in mesenchymal progenitors.

Second, Matrix Mesenchyme maps to locations that are distinct from mesenchymal progenitors, which localize to the periphery of the lung. We also identified interactions between Matrix Mesenchyme and Neurons via PTN-PTPRZ1, and blocking PTN signaling reduces lung innervation. These findings suggest that Matrix Mesenchyme might play a crucial role in innervation of the early embryonic lung. We want to note that while Matrix Mesenchyme shows the highest expression level of *Ptn*, *Ptn* is also expressed in other cell types, including mesenchymal progenitors, precartilage, pericytes, and neurons themselves (**Fig. S18f**). Given that the signals that promote migration of intrinsic neurons into the lung remain unclear^27,53–55^, we postulate that spatial transcriptomics approaches can help fill the gaps by pinpointing the critical ligand-receptor interactions between neuronal progenitors and other cell types.

It is important to note that the gene-expression pattern of Matrix Mesenchyme closely resembles that of Alveolar Fibroblast 2 (AF2), a cell type previously identified in the Mouse Lung Cell Atlas, except that *Mfap5*, a top marker of AF2, is not enriched in Matrix Mesenchyme (**Fig. S17d-e**)^56^. The similarity between Matrix Mesenchyme and AF2 is unexpected, as AF2 is primarily observed postnatally, while our sampling was limited to early embryonic stages^3,32,57,58^. Moreover, the distinct locations of each cell type suggest that Matrix Mesenchyme has a different function than AF2 at this early stage. Future studies are needed to determine how Matrix Mesenchyme matures at later stages and to clarify its relationship with AF2.

Furthermore, our spatial maps reveal that the mesenchymal progenitor subclusters have distinct spatial locations. Specifically, we uncovered three *Wnt2*^high^ mesenchymal progenitor subclusters with distinct spatiotemporal patterns: MP1 is highly proliferative and arises at the lung periphery at *E*11.5, whereas both MP2 and MP3 are less proliferative and appear predominantly from *E*12.5, with MP2 localizing to the lung periphery and MP3 near the vascular endothelium. Based on these features, we reasoned that MP1 is the primary mesenchymal progenitor, although the roles of MP2 and MP3 require further investigation (see **Supplementary Note**).

Moreover, we showed the spatial pseudotime inference for the smooth-muscle differentiation trajectory. Importantly, we observed spatial enrichment between maturing MP1 cells and Smooth Muscle Progenitors, as well as between Smooth Muscle Progenitors and Smooth Muscle (see **Supplementary Note**). These findings lead us to hypothesize that each progenitor subcluster is spatially proximal to the mature state it commits to, as reflected in its gene-expression profile. If validated by orthogonal approaches in the future, incorporating spatial localization could significantly enhance our ability to disentangle the complex and overlapping trajectories of mesenchymal populations^8,9,32^, both in the embryonic lung and in other organs (see **Supplementary Note**).

To link our mesenchymal subclusters with previous studies, we note that MP1 has a similar gene-expression profile as subepithelial mesenchyme, whereas MP2 and Matrix Mesenchyme resemble submesothelial mesenchyme^31^. However, our subclustering in this study resulted in finer resolution with additional spatial context.

Our study also has limitations. Like all other spatial transcriptomics approaches, our spatial mapping cannot avoid capturing transcripts from a mixture of cells for some pixels, even at close-to-single-cell (10 μm) resolution. However, we leveraged the stereotyped anatomy of embryonic lungs and validated our mapping using orthogonal approaches and findings from previous studies. Additionally, all differential gene-expression analyses were conducted using scRNA-seq data rather than spatial transcriptomic data to avoid any confounding issues. Another limitation is that the top markers of Matrix Mesenchyme are not specific, including *Ptn* and *Prrx1*, making it challenging to deconvolve how other cell types contribute to lung innervation via the PTN-PTPRZ1 signaling pathway. The roles of other gene markers driving innervation terms remain to be investigated. As we were preparing this manuscript, a similar spatial mapping of embryonic lungs using the original DBiT-seq protocol was published^59^. However, that work focuses on more mature embryonic stages and lacks the insight we have on the early embryonic stages, likely due to limited resolution (20 µm) and/or detection sensitivity. Our optimized spatial transcriptomic mapping shows a superior efficiency of transcript recovery at high resolution, which allowed us to reveal the early spatiotemporal pattern of epithelial progenitors and identify mesenchymal subpopulations with distinct distributions and developmental roles, providing a valuable resource for studying lung morphogenesis and innervation.

## Acknowledgements

We thank Dr. Wei Wang and the Genomics Core Facility of Princeton University. We also thank members of the Tissue Morphodynamics Group and Chan Lab for helpful discussions and feedback on the manuscript. This work was supported in part by the NIH [HD099030, HL164861, HD111539, HD144311 (to C.M.N.), and HD111537 (to M.M.C.)] and the NSF (2134935 to C.M.N.). P.Z. was supported in part by a Princeton Bioengineering Initiative - Innovators (PBI2), Distinguished Postdoctoral Fellowship. K.G. was supported in part by a postgraduate scholarship-doctoral (PGS-D) from the Natural Sciences and Engineering Research Council of Canada, the Dr. Margaret McWilliams Predoctoral Fellowship from the Canadian Federation of University Women, the Princeton University Procter Fellowship, and an American Heart Association Predoctoral Fellowship.

## Author contributions

P.Z., M.M.C., and C.M.N. conceptualized the study and designed the experiments. P.Z. and K.G. performed the experiments and collected the data. B.K.L. and P.Z. analyzed the results. P.Z., B.K.L., M.M.C. and C.M.N. interpreted the data and wrote the manuscript. All authors provided input on the final manuscript.

## Methods

### Mice

Embryos were isolated from euthanized timed-pregnant mice (*Mus musculus*, C57BL/6J) at *E*11.5 - 13.5 and stored in chilled phosphate-buffered saline (PBS) prior to dissection. Lungs were dissected following standard protocols in chilled PBS supplemented with 10-U/ml penicillin and 10-µg/ml streptomycin (Sigma-Aldrich). All procedures involving animals were approved by Princeton University’s Institutional Animal Care and Use Committee. Mice were housed in an AAALAC-accredited facility in accordance with the NIH Guide for the Care and Use of Laboratory Animals. This study was compliant with all relevant ethical regulations regarding animal research.

### Embedding and sectioning

To embed samples, the dissected lungs were incubated for 1 h on a shaker at 4°C in 0.3 mL each of 20% sucrose, 30% sucrose, and 1:1 OCT:30% sucrose solution with 0.6-μL DEPC, sequentially, before embedding in disposable base molds using dry ice. The embedded lungs were warmed up at -20°C in a cryostat and sectioned into ∼7-μm thickness before being placed in the center of poly-L-lysine-coated glass slides. The cryosections were stored at -80°C before testing. In total, we collected 5 sections from five *E*11.5 embryonic lungs, 4 sections from three *E*12.5 embryonic lungs (2 out of 4 sections are from a single embryonic lung), and 4 sections from four *E*13.5 embryonic lungs.

### Immunostaining

Lungs were dissected and fixed in 4% paraformaldehyde (PFA) in PBS on a shaker at 4°C for 15 min. Lungs were then washed four times for 15 min with 0.5% Triton X-100 in PBS and blocked overnight at 4°C using 5% donkey serum in PBST on a shaker. Samples were then incubated with primary antibodies against PTN (mouse, 1:100, Santa Cruz Biotechnology sc-74443), E-cadherin (rat, 1:400, Invitrogen 131900), and TuJ1 (rabbit, 1:400, R&D Systems MAB1195-SP) for three days at 4°C, followed by Alexa Fluor-conjugated secondary antibodies (1:400; Thermo Fisher Scientific A78946 and A32788, Biotium 20015) and Hoechst (1:1000) for another 2 days at 4°C.

### Function-blocking antibody experiments

Lungs (*Mus musculus*, CD1) were dissected at early *E*11.5 and then cultured as previously described^60^. Briefly, dissected lung explants were positioned on top of a porous membrane (nucleopore polycarbonate track-etch membrane, 8-μm pore size, Whatman), floating on the surface of DMEM/F12 medium supplemented with 50 units/mL of penicillin and streptomycin, 5% fetal bovine serum (FBS, heat inactivated; Atlanta Biologicals), containing either desalted anti-PTN antibody (6-12 µg/mL; Santa Cruz Biotechnology sc-74443) or mouse IgG1 antibody (Mouse IgG1 Isotype Control, Invitrogen, 026100) as a control. Lungs were cultured for 36 h, then fixed in 4% PFA for 15 min at 4°C, followed by immunostaining. We note that the staging of the embryonic lungs and the culture duration are critical for this inhibition experiment.

### Fluorescence in situ hybridization

Adjacent lung sections were used for fluorescence in situ hybridization to validate the spatial transcriptomic mapping. Specifically, we used the RNAscope analysis Multiplex Fluorescent V2 Assay (ACD) protocol. Probes used were *Mus musculus Shh* (314361-C1), *Sox9* (401051-C3), *Sox2* (401041-C2), *Prrx1* (485231-C1), and *Osr1* (496281-C2). Sections were imaged on a spinning disk confocal (BioVision X-Light V2) fitted to an inverted microscope.

### Microfabrication

Our spatial tests used two microfluidic devices – Chip A for loading barcodes A1-A50 and Chip B for loading barcodes B1-B50 in an orthogonal direction, following a design from previous studies^25^. Master molds were microfabricated via standard photolithography on 4-inch silicon wafers. Specifically, a layer of SU8-2025 photoresist (∼25 μm) was spin-coated and patterned using the Heidelberg DWL66+ direct write lithography system prior to developing. The mold was hard baked at 150°C and silanized for 15 min before device fabrication. Chips A and B were fabricated by adding ∼30-g 10:1 (base to curing agent ratio) Sylgard 184 (Dow Corning 4019862) onto the mold and curing for 2 h at 65°C. The inlets and outlets of each device were then punched and cleaned using IPA for 10 min with sonication before use.

### Spatial sequencing

#### Preprocessing

Sections were taken from the freezer and imaged at the desired optical resolution (4×, 10×, or 20× air objective). The sections were fixed using a mixture of 800-μL ice-cold methanol, 15-μL 50-mg/mL DSP in dimethyl sulfoxide, and 1.6-μL diethyl pyrocarbonate for 15 min. The sections were then cleaned by pipetting 1 mL of PBS-RI (998.6-μL PBS with 1.4-μL RNase inhibitor) across the tissue three times and then dried with gentle air flow.

The tissue sections were then blocked for 30 min in 100-μL BSA (10 mg/mL) in PBS at 4°C in a humidified chamber, followed by washing three times with 100-μL 0.5x PBS-RI for 5 min, followed by another two quick washes using 0.5x PBS-RI.

To start spatial barcoding, the first microfluidic device was centered on the tissue. We then took a scan to record the position of the channels on the tissue. We added and loaded 1-mL PBST into the reservoir to permeabilize the tissue section for 20 min. We then removed the remaining

PBST from the reservoir and the inlets/outlets and washed the channels by adding ∼1-mL 0.5x PBS-RI to the reservoir and flowing for about 10 min. We applied vacuum for 5 min to remove any remaining washing buffer from the channels.

#### Reverse transcription

We first prepared the following RT mixture: 29.2-μL RNase-free water, 55-µL 5x Maxima RT buffer, 110-µL PBS-RI, 1.76-µL RNase inhibitor (Enzymatic), 3.52-µL Superase In RNase Inhibitor (Ambion), 13.75-µL 10-mM dNTPs, 27.5-µL Maxima H Minus Reverse Transcriptase, and 6.88-µL 100-µM template switch oligos. We then combined 4.5-µL RT mixture and 0.5 µL of 25-µM Barcode A1-A50, before introducing the barcode mixture into the inlets of the microfluidic device sequentially. We then applied the vacuum to fill all channels with solution and incubated in a humidified chamber at room temperature for 30 min followed by 42°C for 90 min for in-tissue reverse transcription.

Following reverse transcription, we removed the barcode A mixture from the inlets/outlets and cleaned the channels by flushing with 0.5-mL NEB buffer 3.1 with 1% Enzymatics RNase Inhibitor (10 µL for each inlet) for 10 min. We then peeled off Chip A and briefly dipped the slide in a 50-mL falcon tube with RNase-free water and gently dried under air.

#### Ligation

We then placed and centered Chip B on the tissue, imaging the channels to ensure proper placement. After firmly clamping Chip B on the tissue section, we prepared the ligation mixture as follows: 55.39-µL RNase-free water, 27.5-µL 10× T4 ligase buffer, 11.2-µL T4 DNA ligase (400 U/µL), 2.24-µL RNase inhibitor (Enzymatics), 0.71-µL Superase In RNase Inhibitor (Ambion), 5.5-µL 5% Triton-X-100, and 117.9-µL 1× NEB buffer 3.1 with 1% RNase inhibitor (Enzymatics). We combined 4-µL ligation buffer with 1-µL 25-µM barcode B1-B50, before introducing the barcode B buffer into the channels sequentially. We then incubated in a humidified chamber at 37°C for 30 min for in-tissue ligation of barcode Bs. Following ligation, we cleaned the channels by flushing with wash buffer (500-µL PBS, 5-µL 10% Triton X-100, 1.25-µL Superase In RNase Inhibitor) for 10 min. We then peeled off Chip B and dipped the slide into a 50-mL falcon tube with RNase-free water before gently drying under air. We took a final image of the tissue section to enable mapping and alignment of the sequencing results.

#### Lysis and sub-library generation

To collect tissue-derived cDNA, we added 10 or 15 µL of lysis buffer (0.5x PBS, 10-mM Tris pH 8.0, 200-mM NaCl, 50-mM EDTA pH 8.0, 2.2% SDS, 2-mg/mL Proteinase K) for 10-µm and 15-µm tests, respectively, to the spatially barcoded tissue sections. We then incubated the tissue section at 55°C for 2 h in a tightly sealed and humidified chamber. We collected the lysate into a 1.5-mL DNA low-binding centrifuge tube and then added the same amount of extra lysis solution to wash out the reservoir and retrieve as much lysate as possible. The cell lysate was immediately stored at -80°C.

#### Cell lysate purification

To purify cDNA from cell lysates, we first prepared 40-µL Dynabeads^TM^ MyOne^TM^ Streptavidin C1 beads (Thermo Fisher) per sample, following the supplier’s instructions. We then added 7 volumes of DNA Binding Buffer to each volume of cDNA sample before mixing briefly by vortexing. We purified the mixture using the provided Zymo-Spin™ Column in a Collection Tube. After two washes, we eluted the purified cDNA using 100-µL RNase-free water. We then further captured biotin-labelled cDNAs by mixing 100 µL of resuspended Dynabeads MyOne Streptavidin C1 magnetic beads to each lysate at room temperature for 60 min. We then washed the C1 beads twice with 400 µL of B&W buffer (2.5-mM Tris-HCl, pH 8.0, 0.5-M NaCl, 0.25-mM EDTA, pH 8.0) with 5 min rotation after resuspending beads, followed by another wash once with 400-µL 10-mM Tris-HCl with Tween-20 with 5 min rotation after resuspension.

#### Random-priming-assisted second-strand synthesis

The beads with cDNA captured were first resuspended in 200 μL of ExoI mix [170-μL water, 20-μL ExoI buffer, and 10-μL ExoI (NEB, M0568)] and incubated at 37°C for 50 min.

Following ExoI treatment, the beads were washed three times using 400 μL of B&W buffer with 0.05% Tween (2.5-mM Tris-HCl, pH 8.0, 0.5-M NaCl, 0.25-mM EDTA, pH 8.0, 0.1% Tween-20). The bead pellet was then resuspended in 200 μL of 0.1-M NaOH and incubated for 5 min at room temperature. The supernatant was then discarded and the beads were washed three times in 400 μL of B&W buffer with 0.05% Tween.

The random-priming-assisted second-strand synthesis was then carried out on the beads by incubating them in 200 μL of second-strand synthesis mix (2-µL 1-mM dN-SMRT oligo, 5-µL Klenow enzyme, 20-µL 10-mM dNTPs, 40-µL 5x Maxima RT buffer, 133-µL RNase-free water) at 37°C for 1 h. After the reaction, the supernatant was removed, and the beads were washed three times using 400 μL of B&W buffer with 0.05% Tween-20.

#### PCR

The beads were then washed once with 400 µL of 10-mM Tris and 0.1% Tween-20 solution (with resuspension) and once more with 400 µL of RNase-free water (without resuspension). The beads were then resuspended in the 220-µL PCR mix solution (110-µL 2x Kapa HiFi HotStart Master mix, 8.8-µL 10-µM Primers, and 92.4-µL RNase-free water). The PCR cycles were as follows: 95°C for 3 min, followed by 5 cycles at 98°C for 20 s, 65°C for 45 s, and 72°C for 3 min. The beads were then removed, and the remaining supernatant was used to run the second PCR: 95°C for 3 min, followed by 10-15 cycles at 98°C for 20s, 65°C for 20s, and 72°C for 3 min, followed by 72°C extension for 5 min. We then purified cDNA from the PCR product using 0.8x KAPA pure beads following the supplier’s instructions.

#### Sequencing

The sequencing library was then prepared using a Nextera XT DNA Library Preparation Kit by following the supplier’s instructions. The samples were sequenced using Nova-seq SP 300nt Flowcell with 80 bp for read1, and 250 bp for read 3.

### Computational Methods

#### DBiT-seq processing

Read 1 was reformatted to extract UMI, Barcode A, and Barcode B sequences. The reformatted Read 1 and Read 3 were mapped against the mouse genome (GRCh39), demultiplexed, and annotated (Gencode release M31) using the ST pipeline v1.8.1. The resulting gene-expression matrix was then filtered by removing pixels that did not cover tissues.

#### Transcriptomic integration

Preprocessed DBiT-seq replicates were integrated together with the previously published scRNA-seq reference datasets^30–33^ to provide higher resolution of transcriptomic signals and facilitate cell-state annotation of the DBiT-seq datasets. Prior to integration, to eliminate sequencing-depth-based batch effects, we subsampled the reference datasets to an average of 20k reads per cell and normalized all datasets using SCTransformV2^61^. We then integrated the datasets (IntegrateLayers using RPCA and indicating reference datasets), followed by clustering and UMAP calculation (RunUMAP with dims = 1:13). Once completed, the integrated object was visualized via UMAP and cell-state annotation was carried out using expression of canonical marker genes of expected lung-cell states. To verify cell-state identities, integrated annotations were compared against previous annotations per reference dataset.

#### *Sox2*^+^*Sox9*^+^ epithelial cell quantification in scRNA-seq

Cells were counted as *Sox2*^+^*Sox9*^+^ if the normalized and log-scaled raw counts of both *Sox2* and *Sox9* were larger than 0.1 for only cells from the scRNA-seq reference datasets.

#### Differential expression and GO analysis

Using the integrated dataset described above, differential expression analysis was carried out between pairwise cell-state comparisons using the normalized and log-scaled raw counts for only cells from the scRNA-seq reference datasets to avoid technology-driven batch effects.

Differential expression was calculated using the Wilcoxon rank-sum test offered in the scanpy rank_gene_groups function and the log-fold change was calculated as the log2(Group1 + pseudocount) – log2(Group2 + pseudocount) with a pseudocount of 0.01. To calculate enriched gene sets, enriched genes (p<0.05, log fold change > 1.3) were analyzed using Enrichr.

#### Pseudotime analysis

Pseudotime analysis was then performed using the Monocle 3 package^74^. Since the integrated dataset contains multiple distinct trajectories, we selected only cell states related to each trajectory prior to generation of the pseudotime axis and subsequent gene-module analysis. The dataset subsets were selected using primary marker genes and temporal dynamics: the epithelial subclusters for the Epithelial trajectory; Mesenchymal Progenitor 1, Smooth Muscle Progenitor, and Smooth Muscle for the Smooth Muscle trajectory. The previously calculated UMAP, cell-state assignments, and spatial localization maps were used for visualizing pseudotime trajectories. To characterize gene sets involved in each differentiation trajectory, the Monocle 3 package was used to identify gene modules that co-vary along the pseudotime axis. First, genes that vary significantly across the full pseudotime axis were identified (graph_test, neighbor_graph = 4) using the Morans I spatial autocorrelation analysis. Then, genes were grouped into modules using Louvain community clustering based on the cells that express them (find_gene_modules). The resulting gene modules were investigated by scoring the average gene expression within different cell-state categories and individual cells using the scanpy_score_genes function. Additional genes that vary along the pseudotime axis were identified using linear regression analysis using the scipy_linregress function.

#### Neighborhood-enrichment and localization analysis

Neighborhood-enrichment analysis was carried out using the Squidpy^62^ package (gr.nhood_enrichment) to measure the neighborhood enrichment of each cell state per each individual dataset.

