## Supplementary Material for "Mapping embryonic mouse lung development using enhanced spatial transcriptomics"

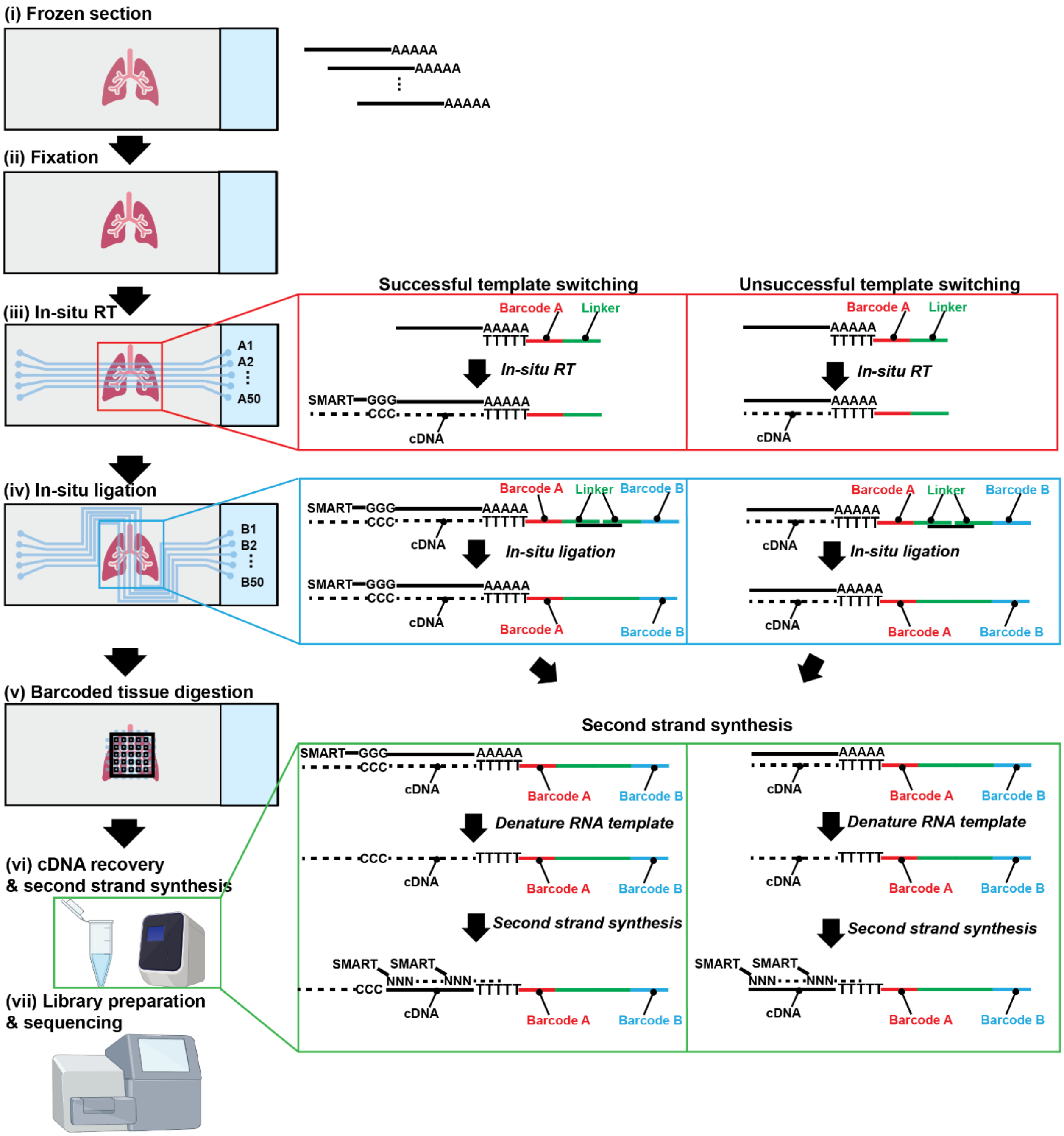


**Fig. S1. Optimized DBiT-seq workflow – related to Fig. 1.** Schematic of steps optimized in a previously published DBiT-seq workflow to enable cellular resolution spatial transcriptomics with high sensitivity.

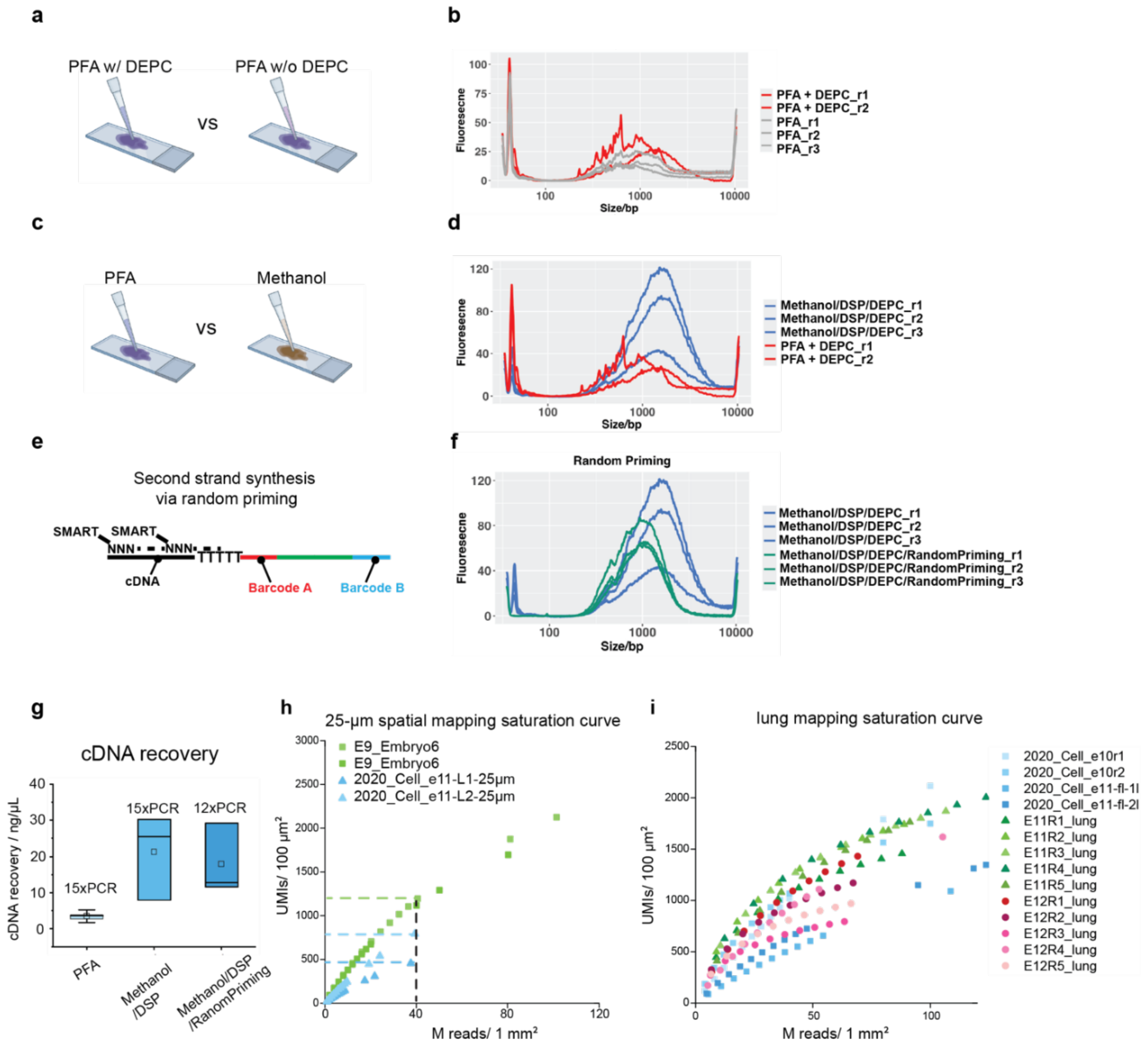


**Fig. S2. Optimized DBiT-seq workflow shows superior cDNA and UMI recovery - related to Fig. 1.** (**a**) Schematic showing comparison between two fixation conditions: PFA with DEPC and PFA without DEPC. (**b**) Graph of cDNA fragment length as a function of fixation condition. (**c**) Schematic showing comparison between two other fixation conditions: PFA with DEPC and methanol/DSP with DEPC. (**d**) Graph of cDNA fragment length as a function of fixation using PFA versus methanol/DSP. (**e**) Schematic showing second-strand synthesis via random priming. (**f**) Graph of cDNA fragment length with or without second-strand synthesis via random priming. (**g**) Graph of cDNA recovery as a function of fixation, with or without an additional random-priming-assisted, second-strand synthesis step. (**h**) Saturation curve of 25-µm optimized DBiT-seq protocol (green) vs original DBiT-seq protocol (blue) when testing whole embryo sections. (**i**) Saturation curve of 10-µm spatial mapping of embryonic lungs using optimized DBiT-seq protocol (green or red) vs 10-µm spatial mapping of whole embryo sections with original DBiT-seq protocol (blue).

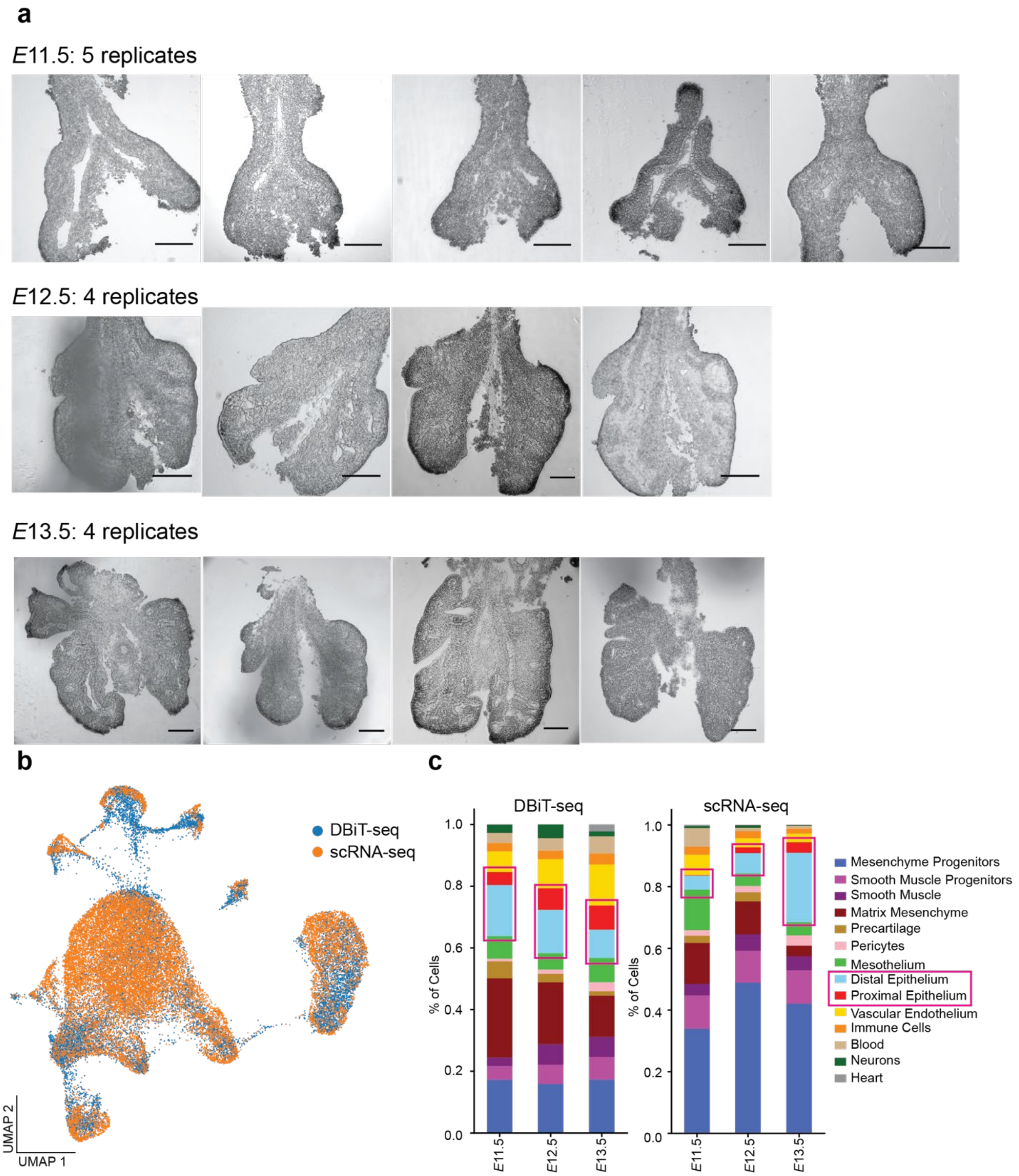


**Fig. S3. Embryonic lung replicates and integration of spatial transcriptomic data with scRNA-seq – related to Fig. 1.** (**a**) Brightfield images of replicates at each stage of development. Scale bars, 200 µm. (**b**) UMAP of integration of spatial transcriptomic data with reference scRNA-seq data. (**c**) Composition of clusters at each stage of development.

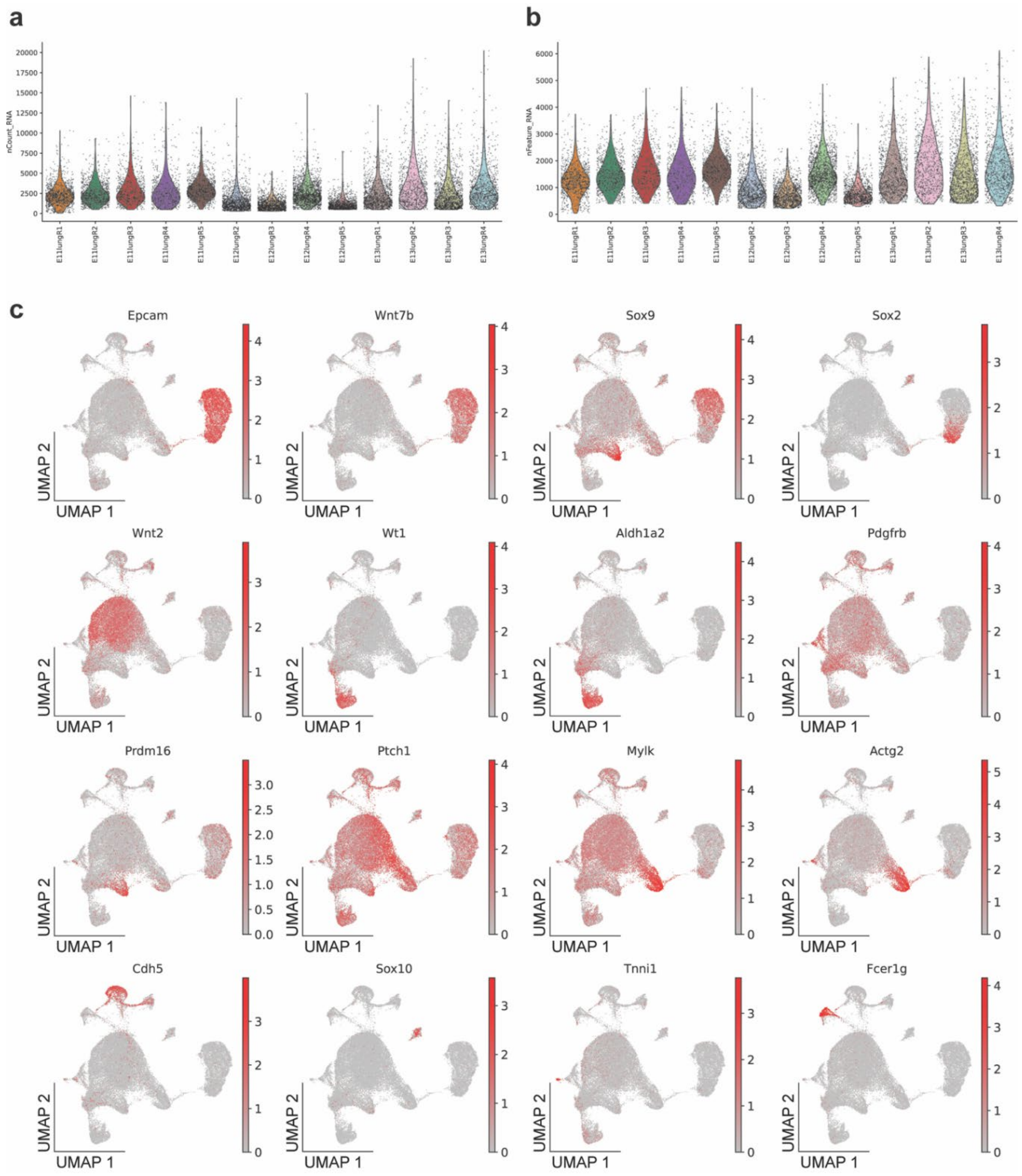


**Fig. S4**. **Spatial transcriptomic data quality control and plots of feature gene markers – related to Fig. 1.** (**a-b**) Violin plots of UMI/pixel and gene/pixel across the different replicates of spatial transcriptomic mapping. (**c**) UMAPs of marker genes for the different cell types in the lung, including *Epcam* and *Wnt7b* (Epithelium), *Sox9* (Precartilage and Distal Epithelium), *Sox2* (Proximal Epithelium), *Wnt2* (Mesenchymal Progenitors), *Wt1* and *Aldh1a2* (Mesothelium), *Pdgfrb* (Pericytes), *Prdm16* (Precartilage), *Ptch1*, *Mylk*, and *Actg2* (Smooth Muscle), *Cdh5* (Vascular Endothelium), *Sox10* (Neurons), Tnni1 (Heart), and *Fcer1g* (Immune Cells).

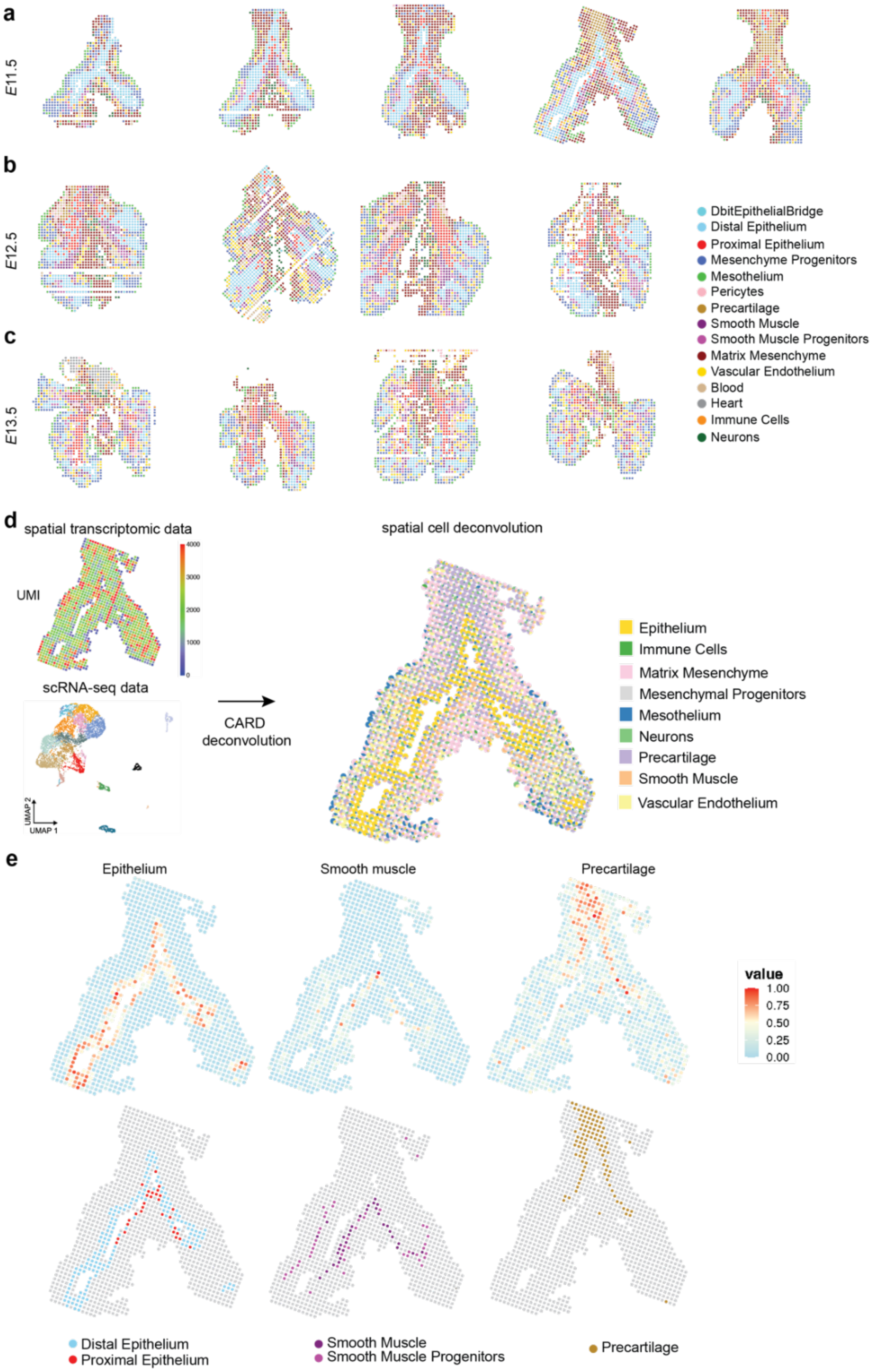


**Fig. S5. Spatial mapping of major cell types across different stages of development – related to Fig. 1.** (**a-c**) Color-coded maps of cells from each cluster, as defined in Fig. 1c, in each replicate at each stage examined. (**d**) Color-coded map of cell identities (right), as inferred by cell deconvolution via CARD using scRNA-seq data (left) as a reference. (**e**) Color-coded maps of epithelium, smooth muscle, and precartilage, as predicted by the deconvoluted data.

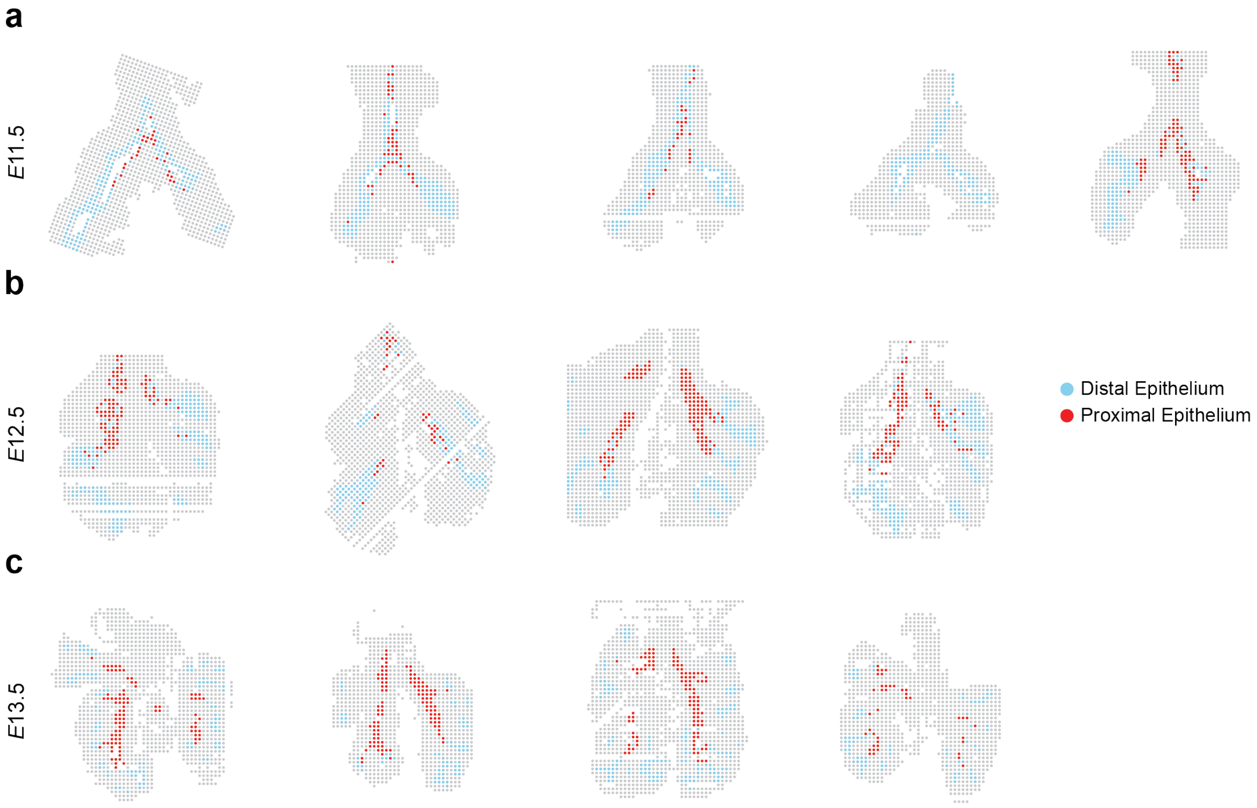


**Fig. S6. Spatial mapping of proximal and distal epithelial cells across different stages of development – related to Fig. 2.** (**a-c**) Color-coded maps of proximal and distal epithelium, in each replicate at each stage examined.

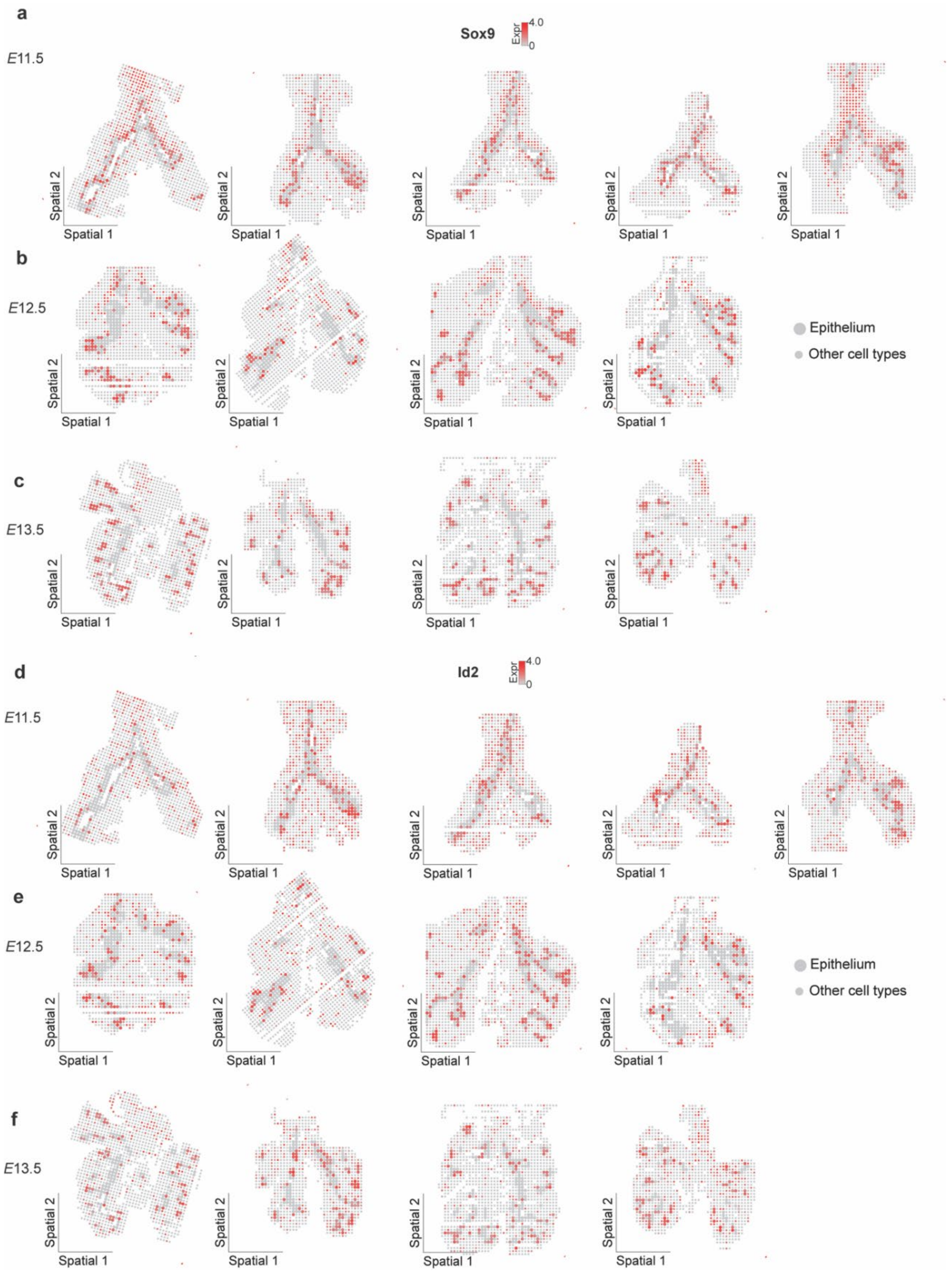


**Fig. S7. Spatial maps of epithelial markers at each stage of lung development – related to Fig. 2.** Color-coded maps of (**a-c**) *Sox9* and (**d-f**) *Id2* in each replicate at each stage examined. For ease of visualization, epithelial cells are highlighted with larger pixels.

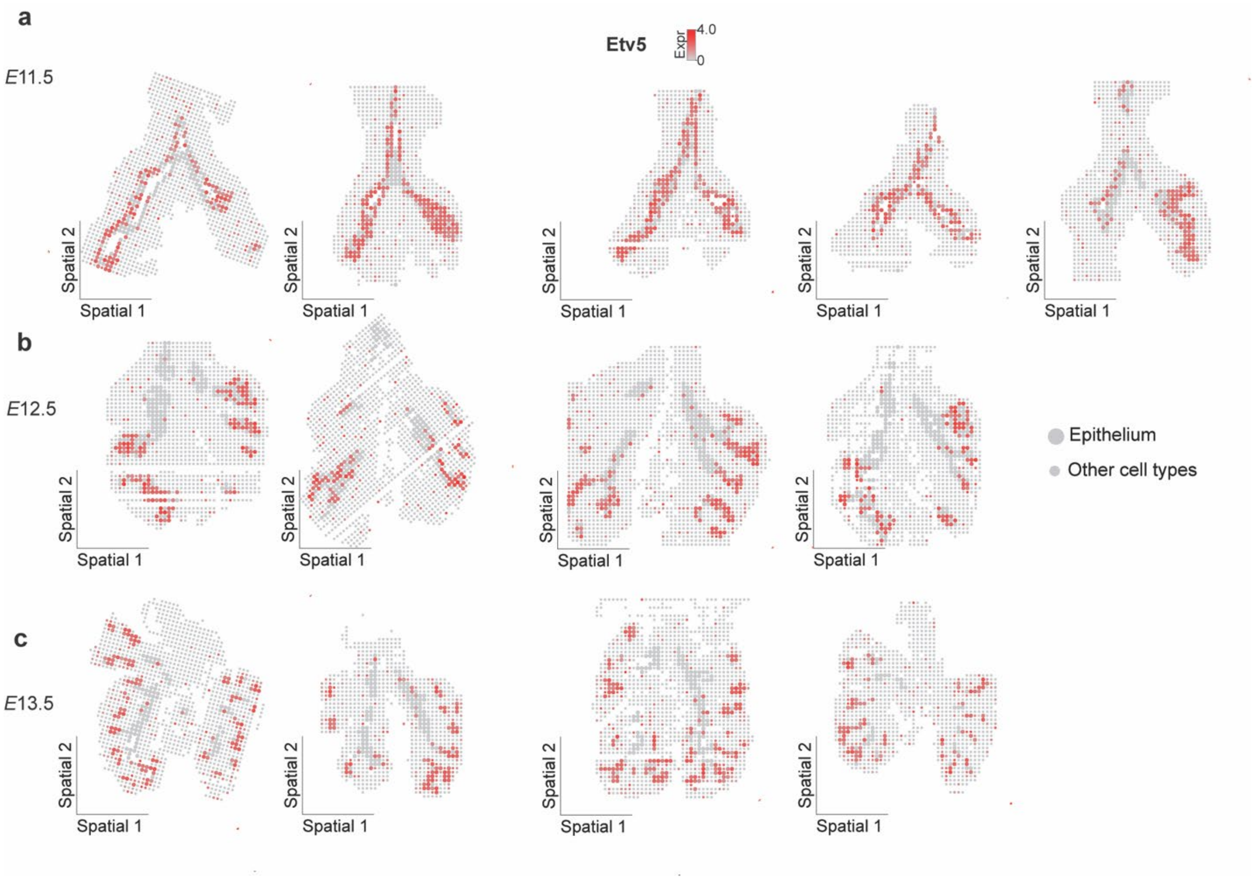


**Fig. S8. Spatial maps of epithelial markers at each stage of lung development – related to Fig. 2.** Color-coded maps of (**a-c**) *Etv5* in each replicate at each stage examined. For ease of visualization, epithelial cells are highlighted with larger pixels.

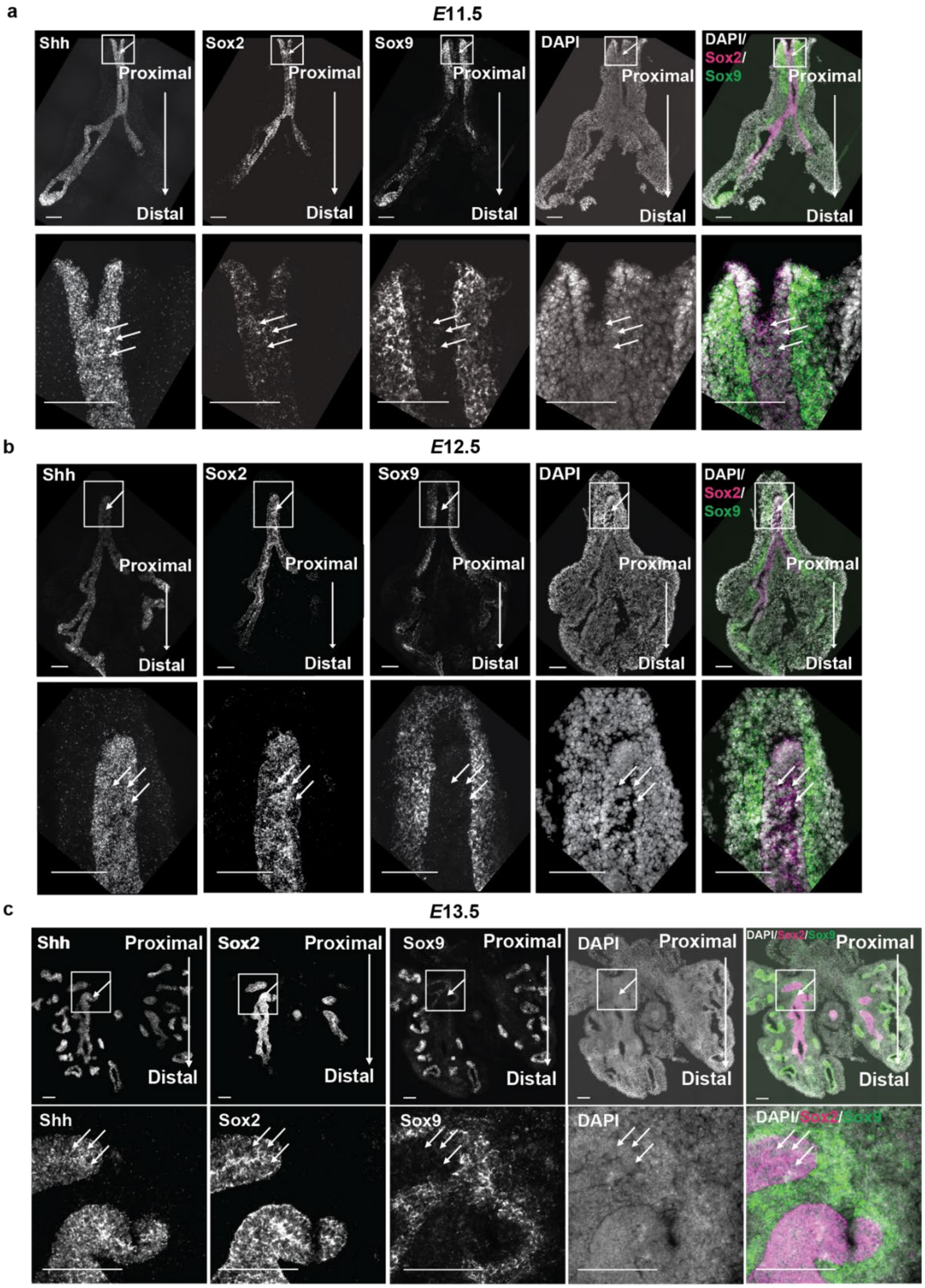


**Fig. S9. Fluorescence in situ hybridization for proximal and distal epithelial markers – related to Fig. 2.** (**a-c**) Top: fluorescence in situ hybridization for *Shh*, *Sox2*, *Sox9*, counterstained with DAPI at each stage examined. Bottom: zoomed-in view of proximal regions boxed in top. Arrows point to the pixels with high *Sox9* signal. Scale bars, 100 µm.

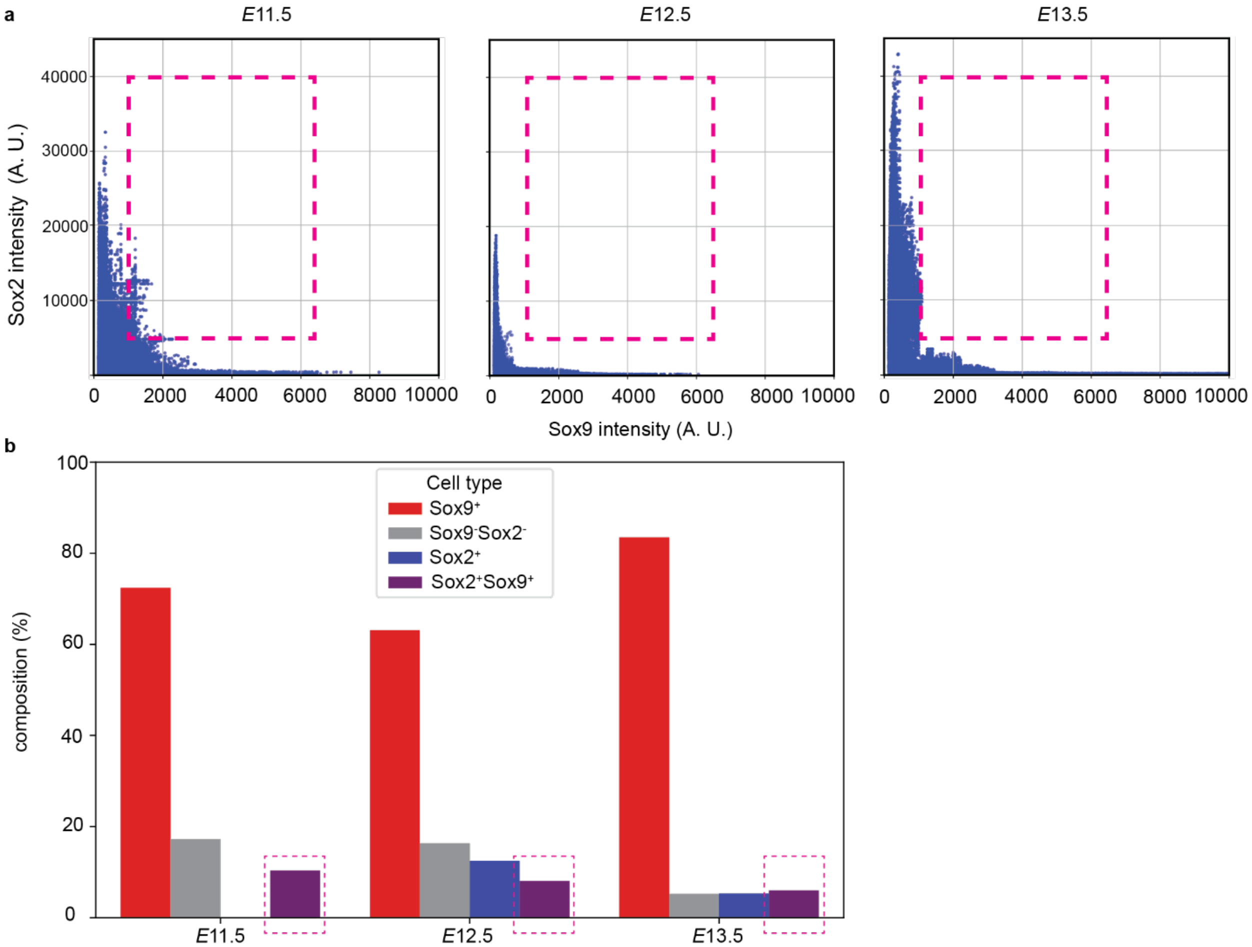


**Fig. S10. Analysis of *Sox2* and *Sox9* expression in the epithelium – related to Fig. 2.** (**a**) Scatter plots of *Sox2* and *Sox9* intensity, from fluorescence in situ hybridization analysis. (**b**) Graph showing quantification of $Sox2^{+}Sox9^{+}$ double-positive epithelial cells in scRNA-seq reference data.

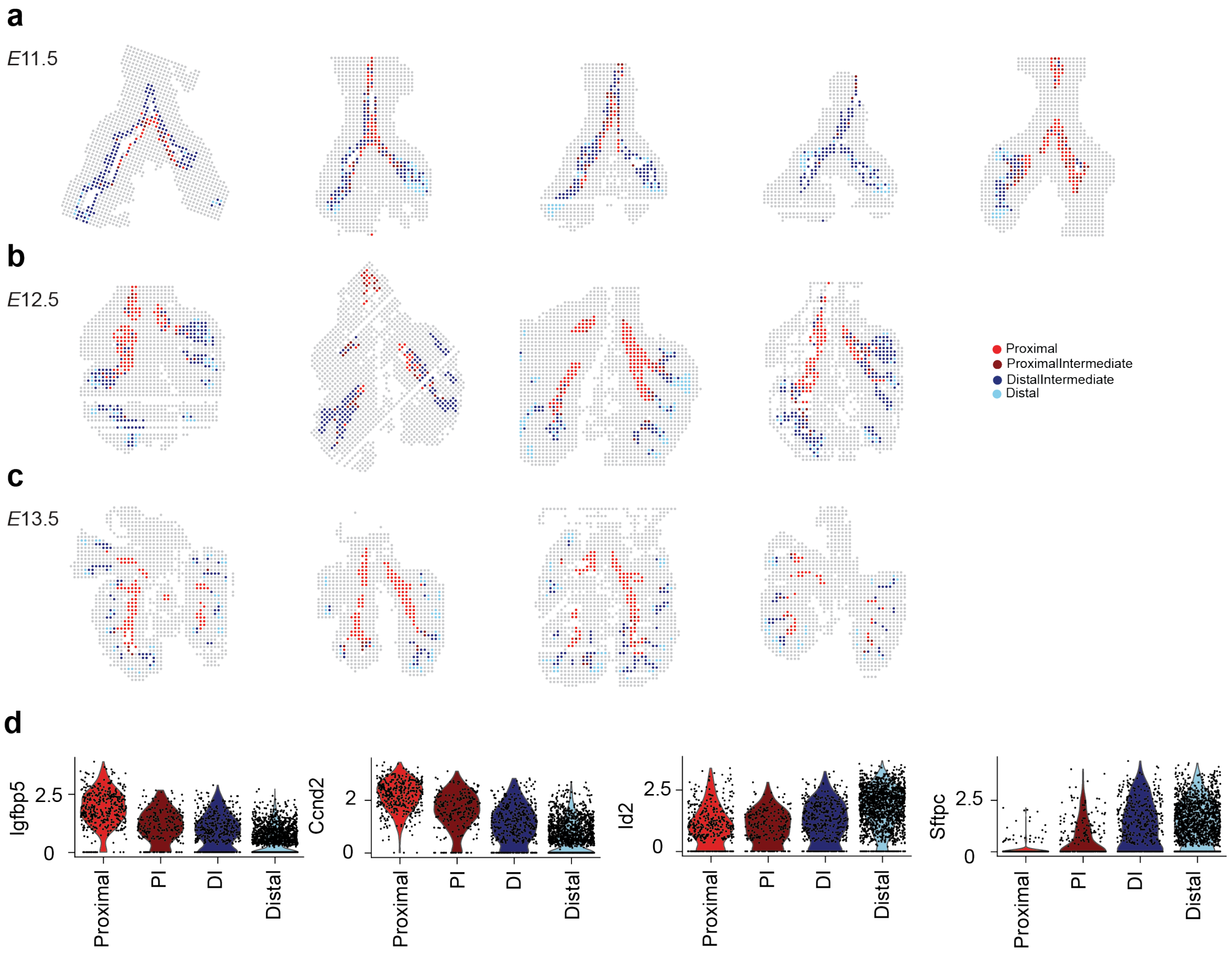


**Fig. S11. Spatial mapping of proximal and distal epithelial cells across different stages of development – related to Fig. 2.** (**a-c**) Color-coded maps of epithelial subclusters, as defined in Fig. 2d, in each replicate at each stage examined. (**d**) Violin plots of additional proximal (*Igfbp5* and *Ccnd2*) and distal (*Id2* and *Sftpc*) markers in each epithelial subcluster.

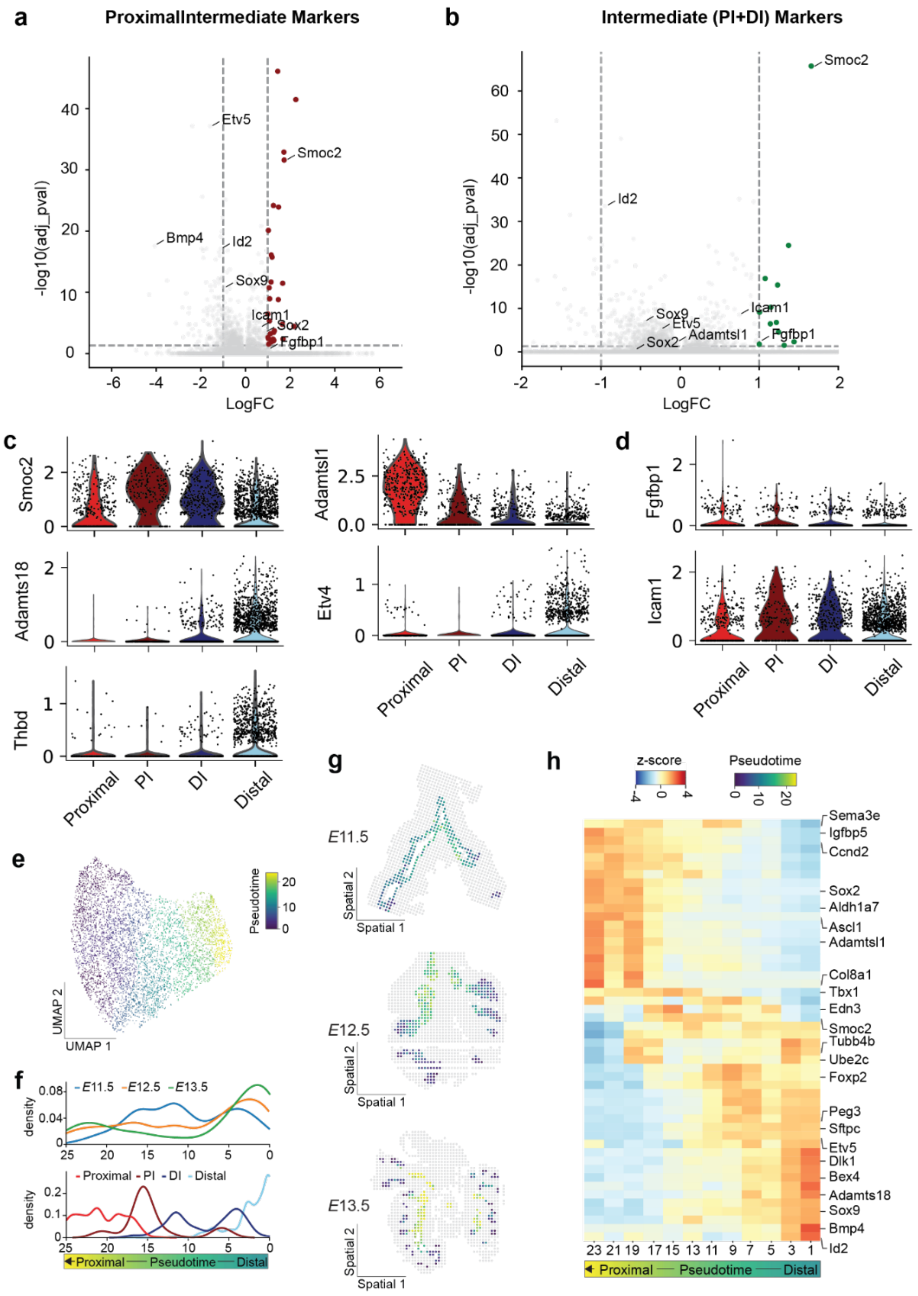


**Fig. S12. Gene markers of epithelial subclusters and pseudotime inference of epithelial differentiation trajectory – related to Fig. 2.** (**a-b**) Volcano plots for differential gene expression between Proximal Intermediate and other epithelial cells, and between merged intermediate and other epithelial cells. (**c**) Violin plots of markers for intermediate (*Smoc2*), Proximal Epithelial (*Adamtsl1*), and Distal Epithelial cells (*Adamts18*, *Etv4*, and *Thbd*). (**d**) Violin plots of *Fgfbp1* and *Icam1*. (**e**) UMAP of epithelial pseudotime inference. (**f**) Graphs of density of cell states along the pseudotime from (e). (**g**) Color-coded maps of spatial pseudotime at each stage of lung development. (**h**) Heatmap of gene markers along the pseudotime trajectory from (e).

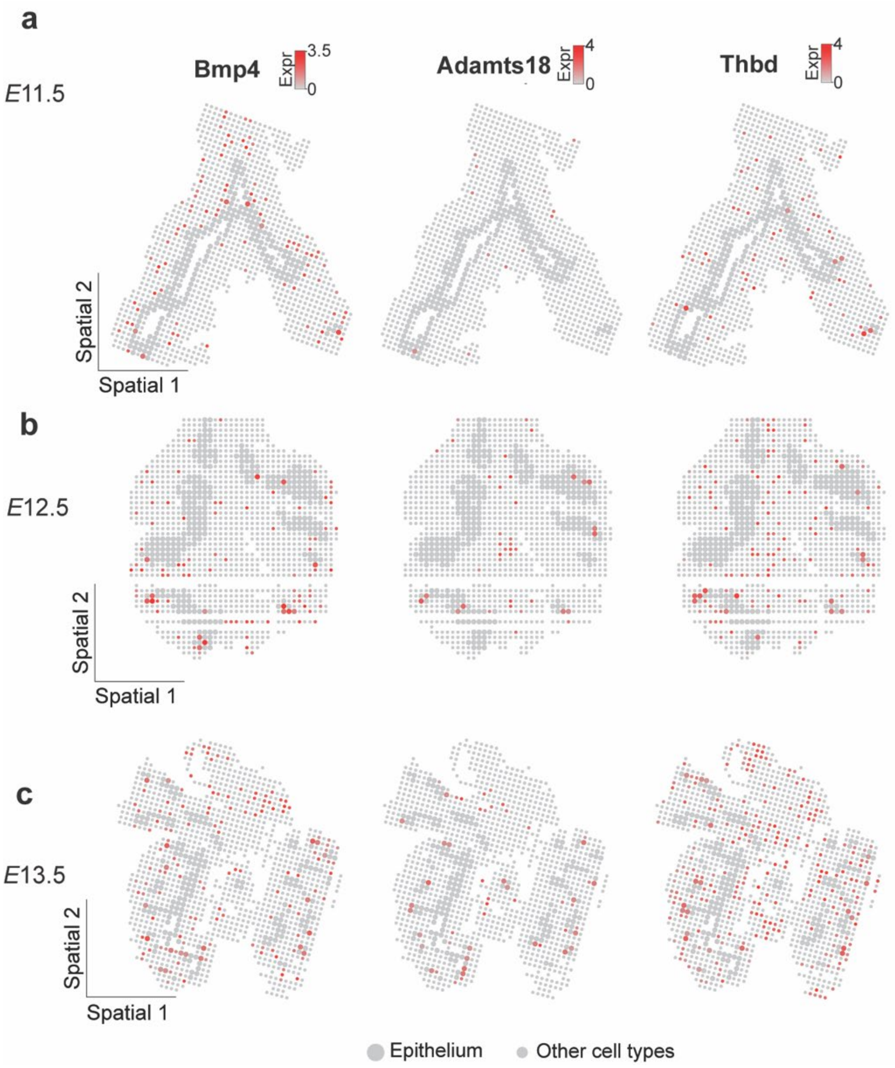


**Fig. S13. Spatial maps of markers for Distal Epithelium – related to Fig. 2.** (**a-c**) Spatial maps of Distal Epithelium-enriched genes *Bmp4*, *Adamts18*, and *Thbd* at each stage examined.

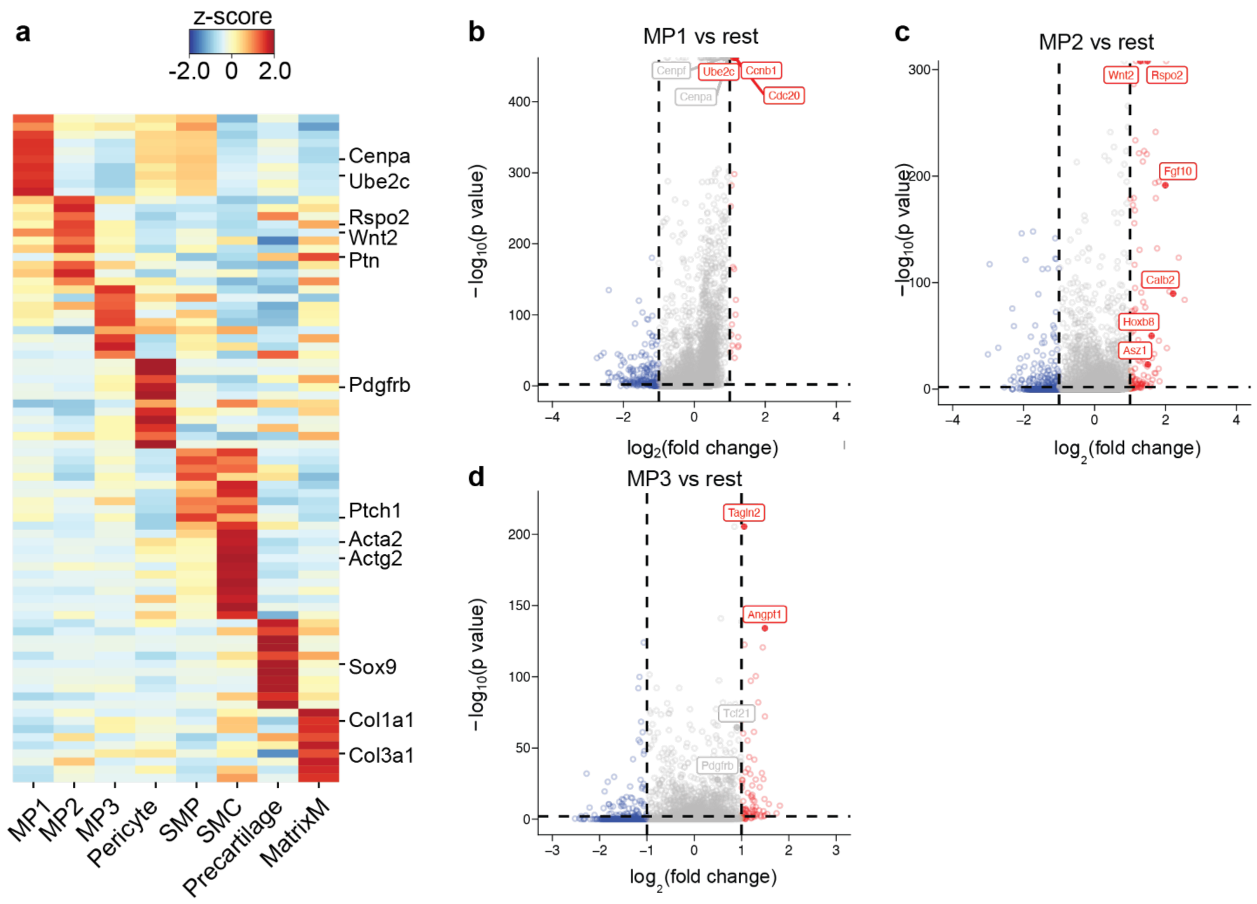


**Fig. S14. Gene markers of mesenchymal subclusters – related to Fig. 3.** (**a**) Heatmap of gene markers for mesenchymal subclusters. (**b-e**) Volcano plots for differential gene-expression analysis of MP1, MP2, and MP3.

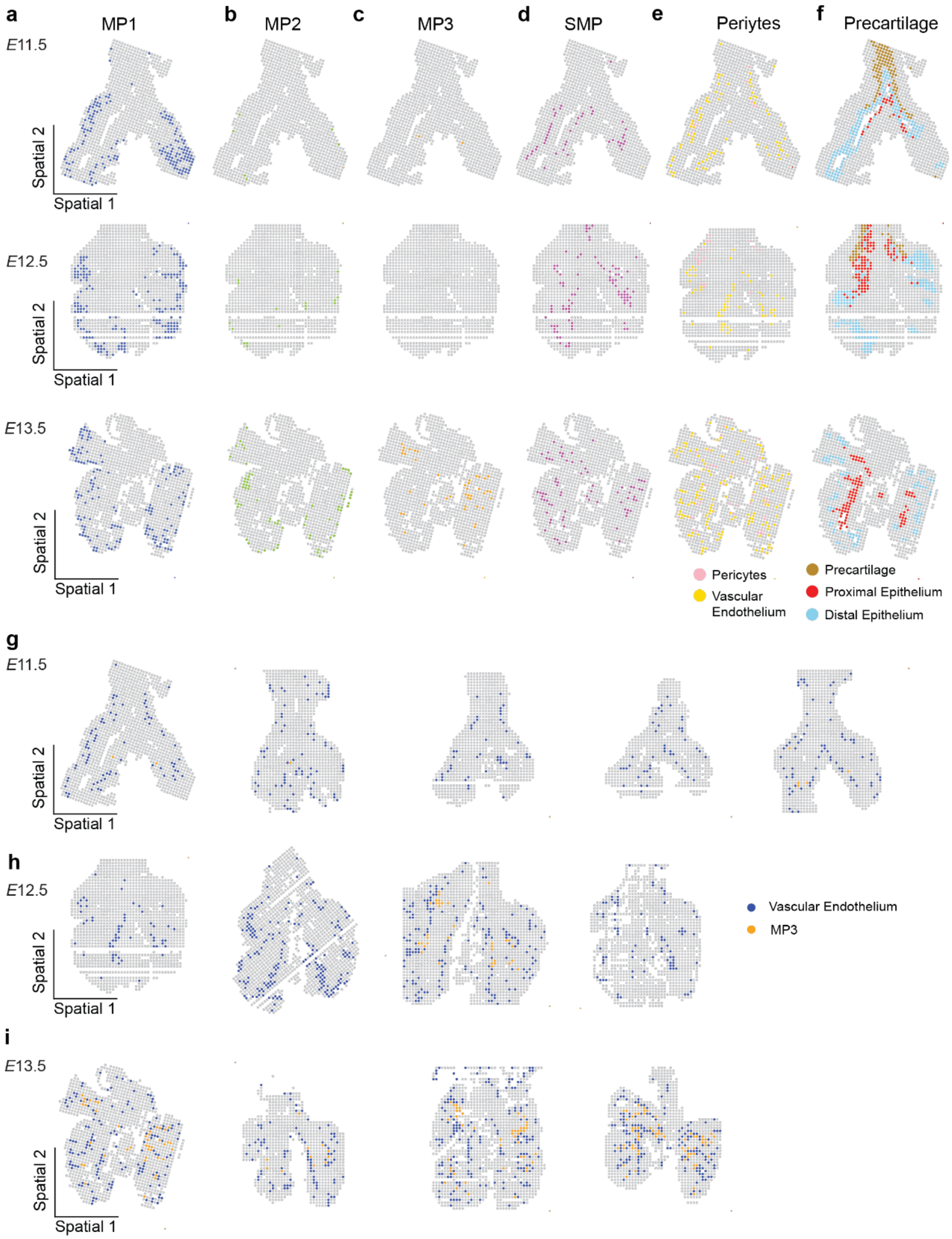


**Fig. S15. Distinct spatial locations of mesenchymal subclusters – related to Fig. 3.** (**a-f**) Color-coded maps of Mesenchymal Progenitor 1 (MP1), Mesenchymal Progenitor 2 (MP2), Mesenchymal Progenitor 3 (MP3), Smooth Muscle Progenitor (SMP), Pericytes, and Precartilage at each stage examined. (**g-i**) Color-coded maps of MP3 with Vascular Endothelium at each stage examined.

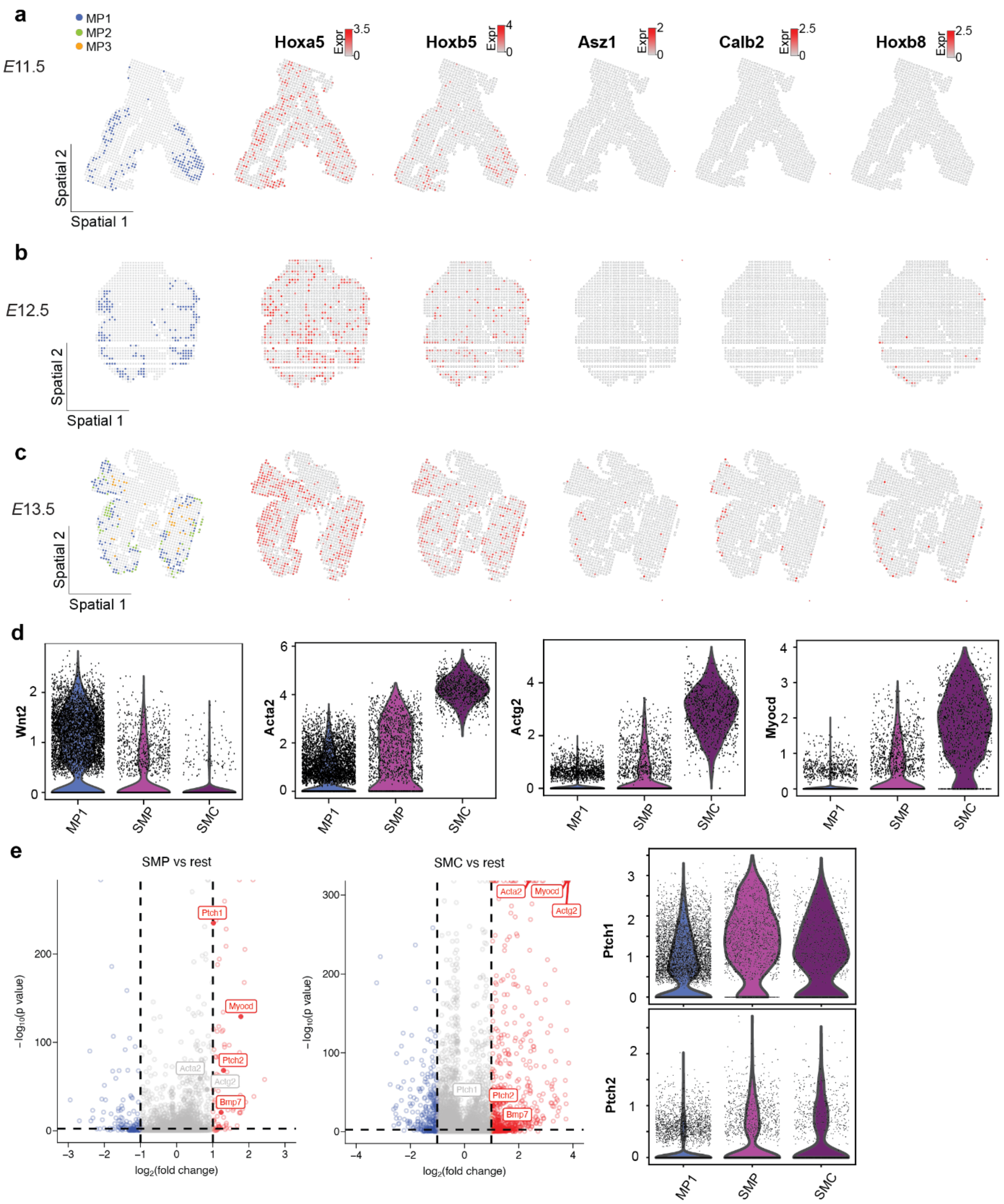


**Fig. S16. Spatial maps of mesenchymal populations – related to Fig. 3.** (**a-c**) Spatial maps of Mesenchymal Progenitor and its top markers, including *Hoxa5*, *Hoxb5*, *Asz1*, *Calb2*, and *Hoxb8*. (**d**) Violin plots of *Wnt2*, and top smooth-muscle-related gene markers for MP1, Smooth Muscle Progenitors (SMP), and Smooth Muscle Cells (SMC). (**e**) Volcano plots for differential gene-expression analysis for Smooth Muscle Progenitors (SMP) and Smooth Muscle Cells (SMC) and violin plots of *Ptch1* and *Ptch2*.

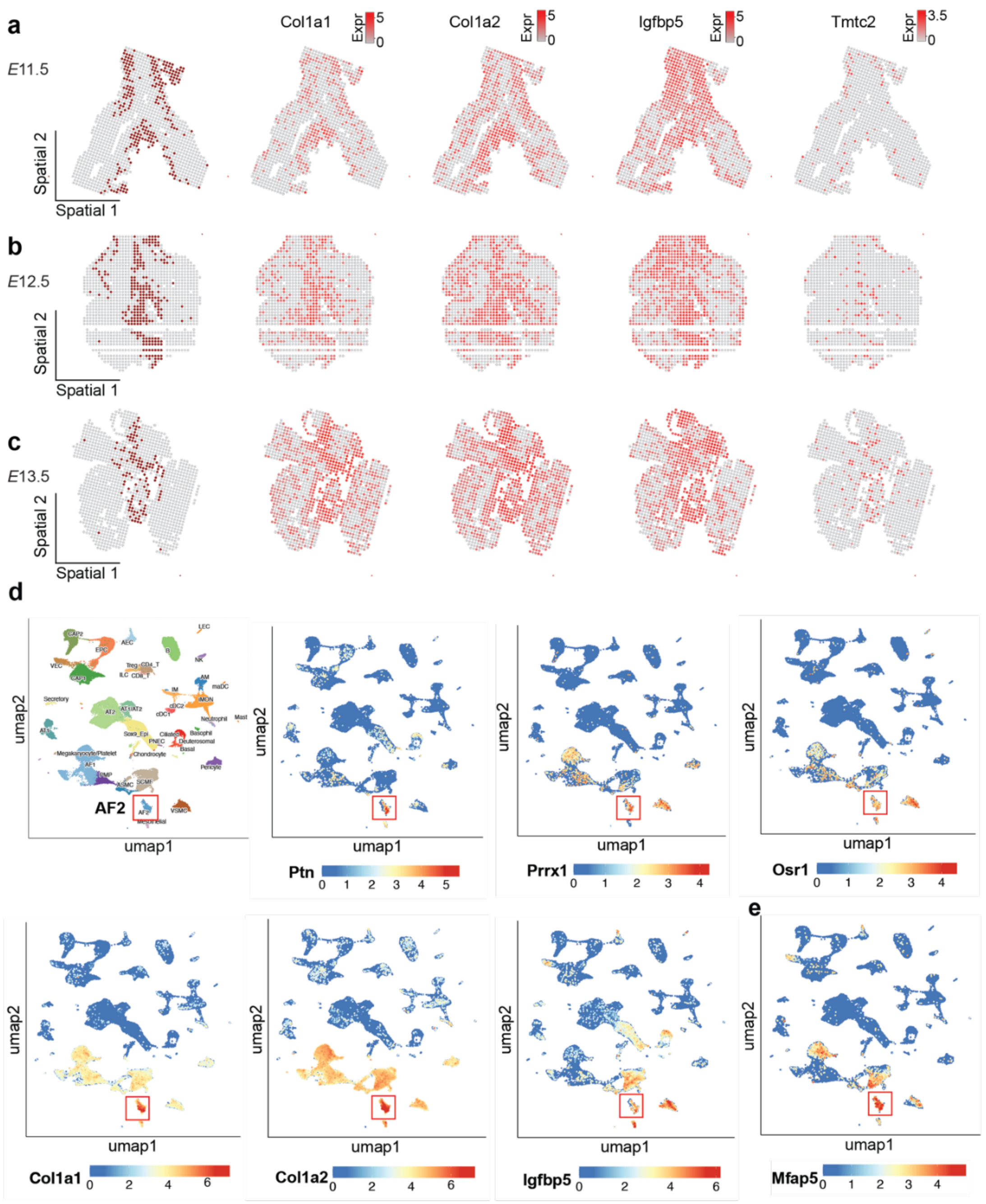


**Fig. S17. Spatial maps of Matrix Mesenchyme, its gene markers, and UMAPs of Alveolar Fibroblast 2 – related to Fig. 4.** (**a-c**) Color-coded spatial maps of Matrix Mesenchyme and its top markers, including *Col1a1*, *Col1a2*, *Igfbp5*, and *Tmtc2*. (**d-e**) UMAP of scRNA-seq reference dataset for mouse lung (Mouse CellCards Multi-Study CellRef 1.0 Atlas), with Alveolar Fibroblast 2 (AF2) highlighted (bottom center) and feature plots of AF2's top markers, including *Ptn*, *Prrx1*, *Osr1*, *Col1a1*, *Col1a2*, *Igfbp5*, and *Mfap5*.

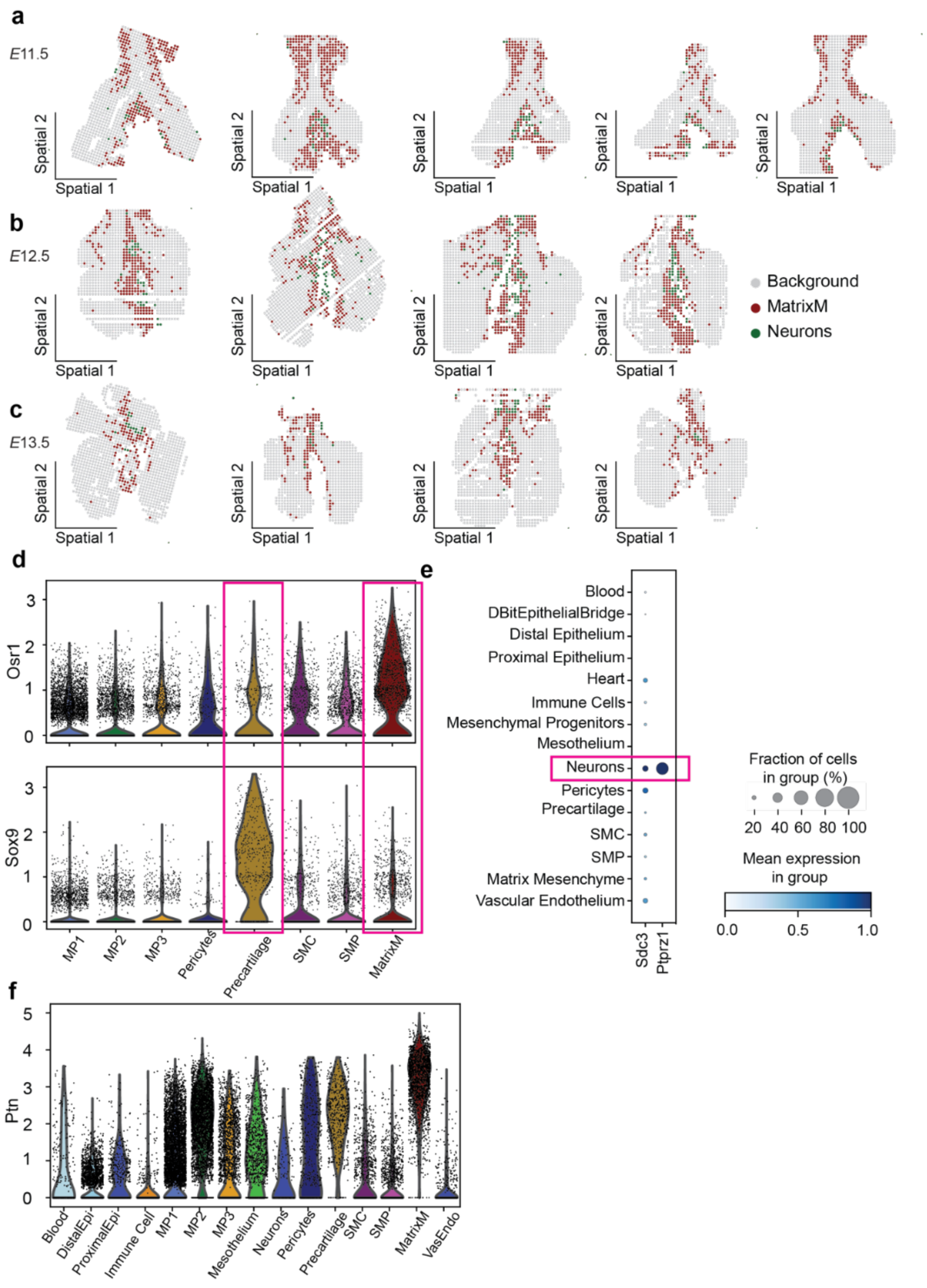


**Fig. S18. Expression of Ptn receptors and spatial localization of Matrix Mesenchyme and Neurons – related to Fig. 4.** (**a-c**) Color-coded maps of Matrix Mesenchyme and Neurons in each replicate at each stage examined. (**d**) Violin plots of *Osr1* and *Sox9* in mesenchymal populations, including Mesenchymal Progenitor 1-3 (MP1-3), Pericytes, Smooth Muscle Progenitors (SMP), Smooth Muscle Cells (SMC), Matrix Mesenchyme (MatrixM), and Precartilage. (**e**) Dot plot of gene expression for Ptn receptors *Sdc3* and *Ptprz1*. (**f**) Violin plot of *Ptn*.

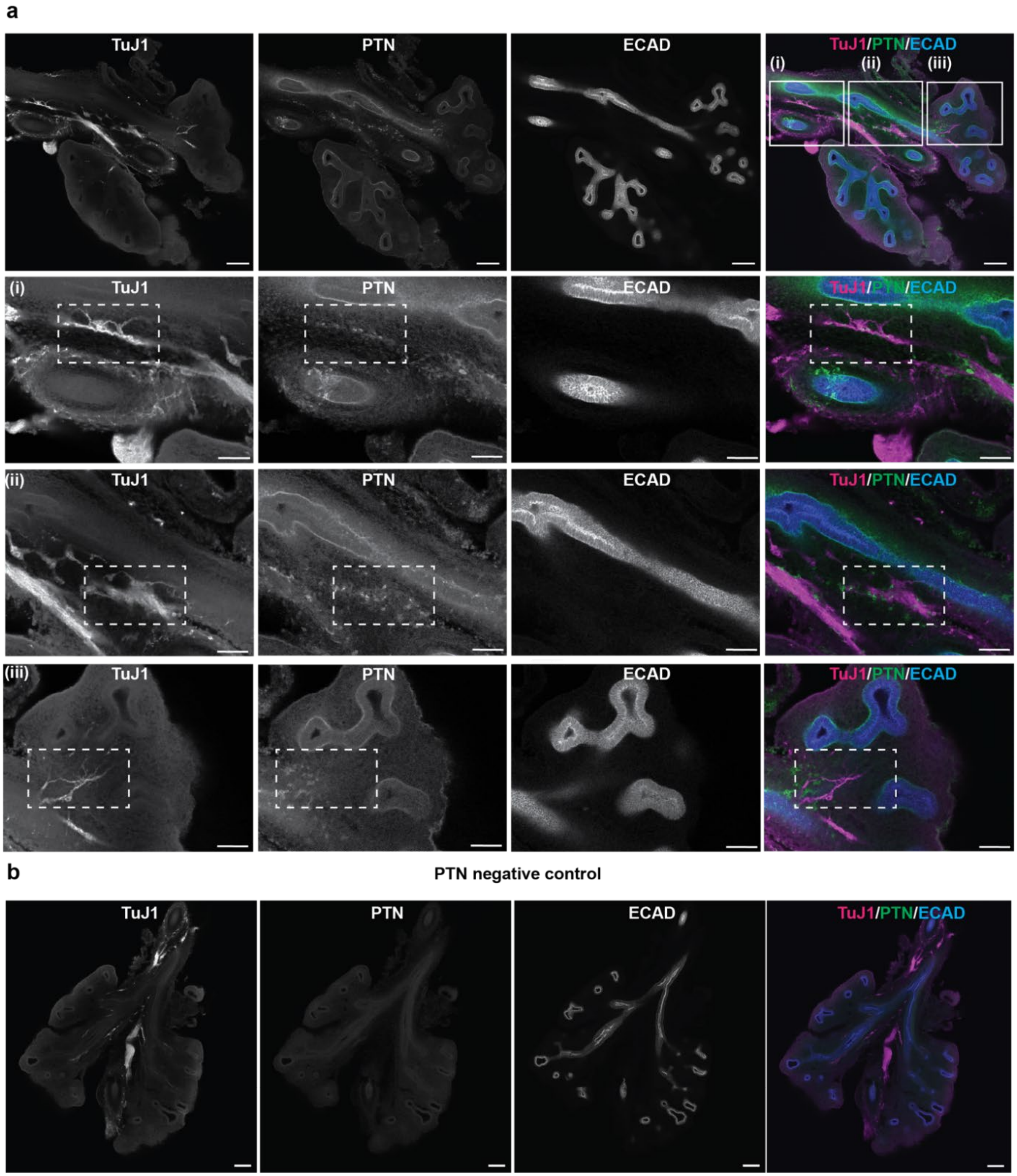


**Fig. S19. Spatial localization of PTN in the embryonic lung– related to Fig. 4.** (**a**) Immunofluorescence staining of the whole lung sections showing TuJ1 (neuronal marker), PTN, and ECAD (epithelial marker). Scale bars, 100 µm. (**i-iii**) Zoomed-in images of TuJ1, PTN and ECAD at three different locations. Scale bars, 50 µm. (**b**) Negative control of immunofluorescence staining for PTN while TuJ1 and ECAD were stained as usual. Scale bars, 100 µm.

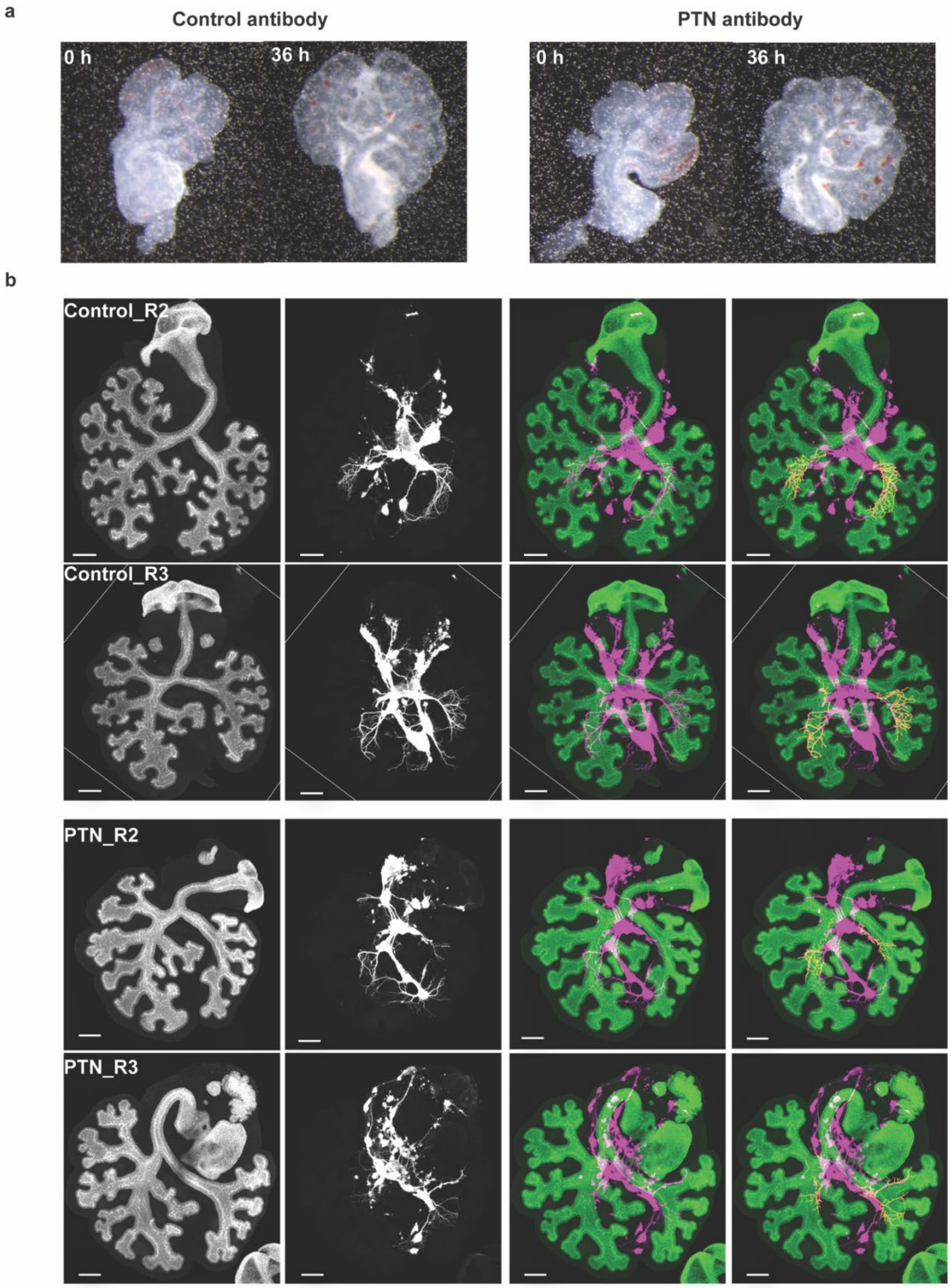


**Fig. S20. Lung explant culture using PTN function-blocking antibody – related to Fig. 4.** (**a**) Ex vivo culture of lung explant using anti-PTN functional blocking or control antibody. (**b**) Immunofluorescence analysis for ECAD (green) and TuJ1 (magenta) in lung explants cultured in the presence of anti-PTN or control antibody for 48 h. Scale bars, 200 μm.

### Supplementary Note

#### Inferred differentiation trajectories for other mature mesenchymal states

We hypothesized that integrating spatial location would allow us to better understand the differentiation trajectory of the pulmonary mesenchyme. Specifically, we assumed that each progenitor subcluster is located adjacent to the mature state to which it commits, as reflected in its gene-expression profile. When mapping pseudotime for the MP1-Smooth Muscle Progenitor-Smooth Muscle differentiation trajectory, we observed a clear spatial enrichment pattern corresponding to cell maturation (**Fig. 3h**). Neighborhood-enrichment analysis among the mesenchymal subclusters confirmed that the Smooth Muscle Progenitor population localizes close to the mature Smooth Muscle population (**Fig. S21a**), although MP1 does not show spatial enrichment with the Smooth Muscle Progenitor. One possible reason for this observation is that MP1, as a primary mesenchymal progenitor source, also contributes to other lineages instead of specifically giving rise to Smooth Muscle Progenitor. Indeed, when we segmented MP1 into three stages along the pseudotime axis, including $MP1_{early}$, $MP1_{middle}$, and $MP1_{late}$, we observed significant enrichment between $MP1_{late}$ and SMP, suggesting the maturing MP1 population becomes spatially proximal to SMP (**Fig. S21b-c**).

In addition to the trajectory of MP1 - Smooth Muscle Progenitor - Smooth Muscle, we also sought to investigate the differentiation trajectories for other mature states. Pairwise differential gene-expression analysis showed that MP2 expresses a higher level of *Rspo2*, *Fgf10*, and genes that are highly enriched in Matrix Mesenchyme including *Ptn, Igfbp5, Dcn* (**Fig. S21d**). Pseudotime analysis also revealed that MP2 is an intermediate state in the Matrix Mesenchyme (**Fig. S21e**). However, we observed that MP2 emerges at a later timepoint than Matrix Mesenchyme (**Fig. S15b, Fig. S18a-c**), suggesting that MP2 is a distinct lineage from the Matrix

Mesenchyme. Moreover, neighborhood-enrichment analysis also revealed that MP2 is negatively enriched with Matrix Mesenchyme, further showing that MP2 is unlikely to be the progenitor for Matrix Mesenchyme (**Fig. S21a**). We noticed that AlveolarFibroblast2, a cell type previously identified in the Mouse Lung Cell Atlas, is also highly enriched in similar genes as MP2 and Matrix Mesenchyme (**Fig. S17d-e**) and is located in the distal alveolar region. As AlveolarFibroblast2 appears much later in lung development and is located similarly to MP2, we therefore postulate that MP2 might be progenitors for AlveolarFibroblast2. Nevertheless, the exact lineage relationship between different progenitor states and mature cell types requires further investigation.

Moreover, we also found that MP3 is proximal to the Vascular Endothelium, as shown in the neighborhood enrichment analysis (**Fig. S15g-i, and Fig. S21a**). In pairwise differential gene-expression analysis, MP3 expresses a high level of *Angpt1, Pdgfrb,* and other Pericyte/Vascular smooth muscle (VSMC)-related genes such as *Col6a1*, *Col6a2*, *Adcy2*, *Nrg2*, *Eln* (**Fig. S14d, Fig. S21f-h**). Taken together, MP3 is likely a progenitor for pericytes and/or vascular smooth muscle.

Collectively, integrating spatial neighborhood enrichment and gene-expression profiles suggests that MP3 and Smooth Muscle Progenitors exhibit early lineage commitments toward Pericyte/Vascular Smooth Muscle and Smooth Muscle, respectively (**Fig. S21i**). The spatiotemporal pattern of MP2, together with its gene-expression profile, suggest that MP2 could be progenitors for AF2, although the exact lineage relationship needs further exploration (**Fig. S21i**). For Matrix Mesenchyme, there might be a mesenchymal state at earlier time points, for example *E*10.5, serving as an intermediate state before differentiation (**Fig. S21i**).

Although our proposed mesenchymal differentiation trajectories are purely inferred from gene expression and spatial neighborhood enrichment, they are partly supported by previous lineage-tracing experiments. Specifically, *Pdgfrb*$^{+}$ progenitors at E12.5[1] and *Tcf21*+ progenitors at E11.5-13.5[2] contribute to pericytes and VSMC, consistent with our placement of MP3 ahead of both populations, as *Pdgfrb* and *Tcf21* are both relatively enriched in MP3 compared to MP1 (**Fig. S21g-h**). Moreover, *Acta2*$^{+}$ progenitors at *E*12.5 give rise to airway smooth muscle, matching our placement of Smooth Muscle Progenitors (SMP) ahead of Smooth Muscle (SM)[3]. To fully validate the proposed differentiation trajectories, future lineage-tracing experiments with precise spatial and temporal resolution will be essential.

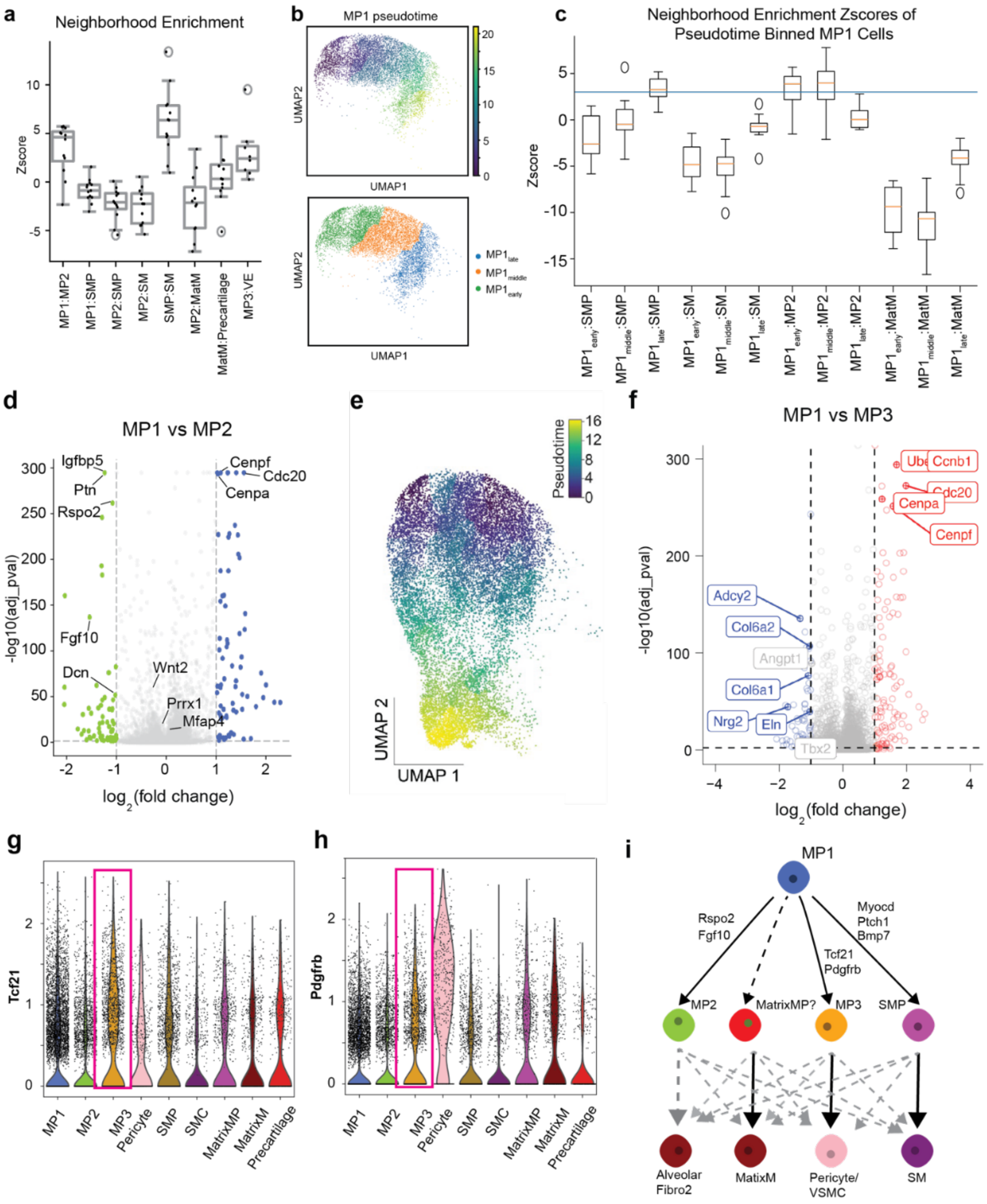


**Fig. S21. Inferred differentiation trajectories for other mature mesenchymal states – related to Supplement Note.** (**a**) Z-score plot for neighborhood-enrichment analysis of mesenchymal subclusters. (**b**) UMAP of segmented MP1, including MP1_early, MP1_middle, and MP1_late, together with color-coded pseudotime inference. (**c**) Z-score plot for neighborhood-enrichment analysis of segmented MP1 with other mesenchymal populations. (**d**) Volcano plot for pairwise differential gene-expression analysis between Mesenchymal Progenitor 1 (MP1) and Mesenchymal Progenitor 2 (MP2). (**e**) Pseudotime inference from MP1, through MP2, to Matrix Mesenchyme. (**f**) Volcano plot for pairwise differential gene-expression analysis between MP1 and Mesenchymal Progenitor 3 (MP3). (**g-h**) Violin plots of *Tcf21* and *Pdgfrb*. (**i**) Schematic of the proposed mesenchymal differentiation trajectory.
